# Deletion of *Crtc1* leads to hippocampal neuroenergetic impairments associated with depressive-like behavior

**DOI:** 10.1101/2020.11.05.370221

**Authors:** Antoine Cherix, Carole Poitry-Yamate, Bernard Lanz, Olivia Zanoletti, Jocelyn Grosse, Carmen Sandi, Rolf Gruetter, Jean-René Cardinaux

**Affiliations:** Laboratory for Functional and Metabolic Imaging (LIFMET), École Polytechnique Fédérale de Lausanne (EPFL), Lausanne, Switzerland; Center for Psychiatric Neuroscience and Service of Child and Adolescent Psychiatry, Department of Psychiatry, Lausanne University Hospital and University of Lausanne, Prilly-Lausanne, Switzerland; Animal Imaging and Technology (AIT), Center for Biomedical Imaging (CIBM), École Polytechnique Fédérale de Lausanne (EPFL), Lausanne, Switzerland; Laboratory of Behavioral Genetics, Brain and Mind Institute, School of Life Sciences, École Polytechnique Fédérale de Lausanne (EPFL), Lausanne, Switzerland

**Keywords:** Depression, Metabolic syndrome, Neuroimaging, Brain metabolism, Hippocampus, Preclinical model

## Abstract

Mood disorders (MD) are a major burden on society as their biology remains poorly understood, challenging both diagnosis and therapy. Among many observed biological dysfunctions, homeostatic dysregulation, such as metabolic syndrome (MeS), shows considerable comorbidity with MD. Recently, CREB-regulated transcription coactivator 1 (CRTC1), a regulator of brain metabolism, was proposed as a promising factor to understand this relationship. Searching for imaging biomarkers and associating them with pathophysiological mechanisms using preclinical models, can provide significant insight into these complex psychiatric diseases and help the development of personalized healthcare. Here, we used neuroimaging technologies to show that deletion of *Crtc1* in mice leads to an imaging fingerprint of hippocampal metabolic impairment related to depressive-like behavior. By identifying the underlying molecular/physiological origin, we could assign an energy-boosting mood-stabilizing treatment, ebselen, which rescued behavior and neuroimaging markers. Finally, our results point towards the GABAergic system as a potential therapeutic target for behavioral dysfunctions related to metabolic disorders. This study provides new insights on *Crtc1’s* and MeS’s relationship to MD and establishes depression-related markers with clinical potential.

## Introduction

Mood disorders (MD) are among the leading causes of disability worldwide^1,2^. The difficulty in defining appropriate treatments relates to the fact that these complex, dynamic and multifactorial psychiatric diseases are poorly understood^3^. The way these diseases are approached complicates the identification of therapeutic targets: diagnosis is currently based on subjective signs and symptoms, rather than on objective biological or chemical measurements. In practice, there is an urgency to establish reliable biomarkers within a framework of personalized treatment approaches^4,5^. Neuroimaging techniques, such as magnetic resonance imaging (MRI), spectroscopy (MRS) and positron emission tomography (PET) are promising tools to achieve this goal by providing brain-specific information^6^. Nevertheless, development and validation of neuroimaging markers for psychiatry require prior understanding of their underlying pathophysiological origin and the genetic and environmental factors linking these markers to behavioral deficit^6^. Among many potential etiological factors that have been identified, metabolic syndrome (MeS), i.e. a combination of obesity, dyslipidemia, insulin resistance, and hypertension, has gained significant attention due to its high co-occurrence with MD^7–10^. However, the mechanisms and the causality relationship between peripheral metabolic alterations and dysfunction of the central mood regulation, and how this translates to *in vivo* brain measurements, remain to be fully elucidated.

The CREB Regulated Transcription Coactivator 1 (*CRTC1*) gene has emerged as a promising target to study how features of MeS can lead to behavioral impairments. Several studies have identified a relationship between *CRTC1* polymorphisms and psychiatric disorders, with focus on obesity parameters^11–14^ and stress^15^. Through its enhancement of CREB transcriptional activity and because of its ability to sense both Ca^2+^ and cAMP second messengers in neurons, CRTC1 has been established as a key regulator of brain function and metabolism^16,17^. CRTC1 is involved in synaptic plasticity and memory formation^18–20^ and participates in the regulation of energy and mood balance^21,22^. Importantly, CRTC1 has been implicated in rodent depressive-like behavior^23^, which can be triggered by excessive CRTC1 phosphorylation and cytoplasmic sequestration as a response to chronic stress^24^. Thus, the *Crtc1* knock-out (*Crtc1*^-/-^) mouse was shown to be a useful model to study the pathways and mechanisms linking metabolic diseases with depression^21,22,25^ and to understand associated resistance to classic antidepressants, in particular to fluoxetine^26,27^.

Here, using state-of-the-art preclinical neuroimaging technologies, we sought to identify fingerprints of brain metabolic disturbances in *Crtc1*^-/-^ mice and to explore its mechanistic relationship with behavioral dysfunctions and MeS. By combining MRS, MRI and PET, we found that deletion of *Crtc1* in mice uncovers hippocampal neuroenergetic markers that are associated with depressive-like behavior. By deciphering the pathophysiological mechanisms associated with these brain markers and behavior, we were able to select a targeted treatment, which reversed the pathological phenotype. Our results highlight new mechanisms linking *Crtc1* and MeS with the development of depressive-like behavior, bringing to the forefront associated preclinical neuroimaging markers with clinical potential, and identification of a compatible mood-stabilizer with therapeutic capacity.

## Results

### Deletion of *Crtc1* is associated with a neuroimaging fingerprint of reduced hippocampal neuroenergetics

We first determined whether deletion of *Crtc1* in mice has measurable metabolic consequences in the brain using proton MRS (^1^H-MRS) and MRI. Animals were scanned at an early age (6 weeks) in a 14.1 Tesla scanner (Fig.1a) to acquire MRI whole brain anatomical images and ^1^H-MRS spectra of the dorsal hippocampus (DH) and cingulate prefrontal cortex (PFC). When comparing the neurochemical profiles of *Crtc1*^-/-^ mice as compared to their wild-type (*Crtc1*^+/+^) littermates (Fig.1b-c), hippocampal neuroenergetic alterations were noted, including a reduced ratio of phosphocreatine relative to creatine (PCr/Cr; *P*=0.04) and increased level of lactate (Lac; *P*=0.02). Subsequently, to evaluate the PCr to Cr ratio measured *in vivo*, high-resolution ^1^H- and ^31^P-NMR of hippocampal metabolite extracts (Fig.1d-g) was performed in another group of mice to further assess the drop in PCr (Fig.1e; *P*=0.04). In addition, an increase in inorganic phosphate was observed (P_i_; *P*=0.03), in line with higher PCr hydrolysis, while ATP levels and the NADH/NAD^+^ ratio were similar in both groups (Fig.1g, n.s.). Interestingly, the neurochemical profile of PFC (Fig.S1a,b) did not indicate neuroenergetic alterations, but an increase in total choline (tCho; *P*=0.0006), i.e. glycerophosphorylcholine (GPC) and phosphocholine (PCho), in *Crtc1*^-/-^ mice. This rise in phospholipid-related metabolites coincided with bigger prefrontal volume (Fig.S1c), as measured from MRI images, suggesting potential prefrontal inflammation. These distinct observations between PFC and DH could not be attributed to differences in *Crtc1* brain regional expression as relative mRNA content was comparable between both regions in the wild-type mice (Fig.S1d). Taken together, these results indicate that *Crtc1* deletion affects hippocampal energy metabolism and prefrontal integrity, producing a measurable fingerprint using neuroimaging modalities.

**Figure 1:**
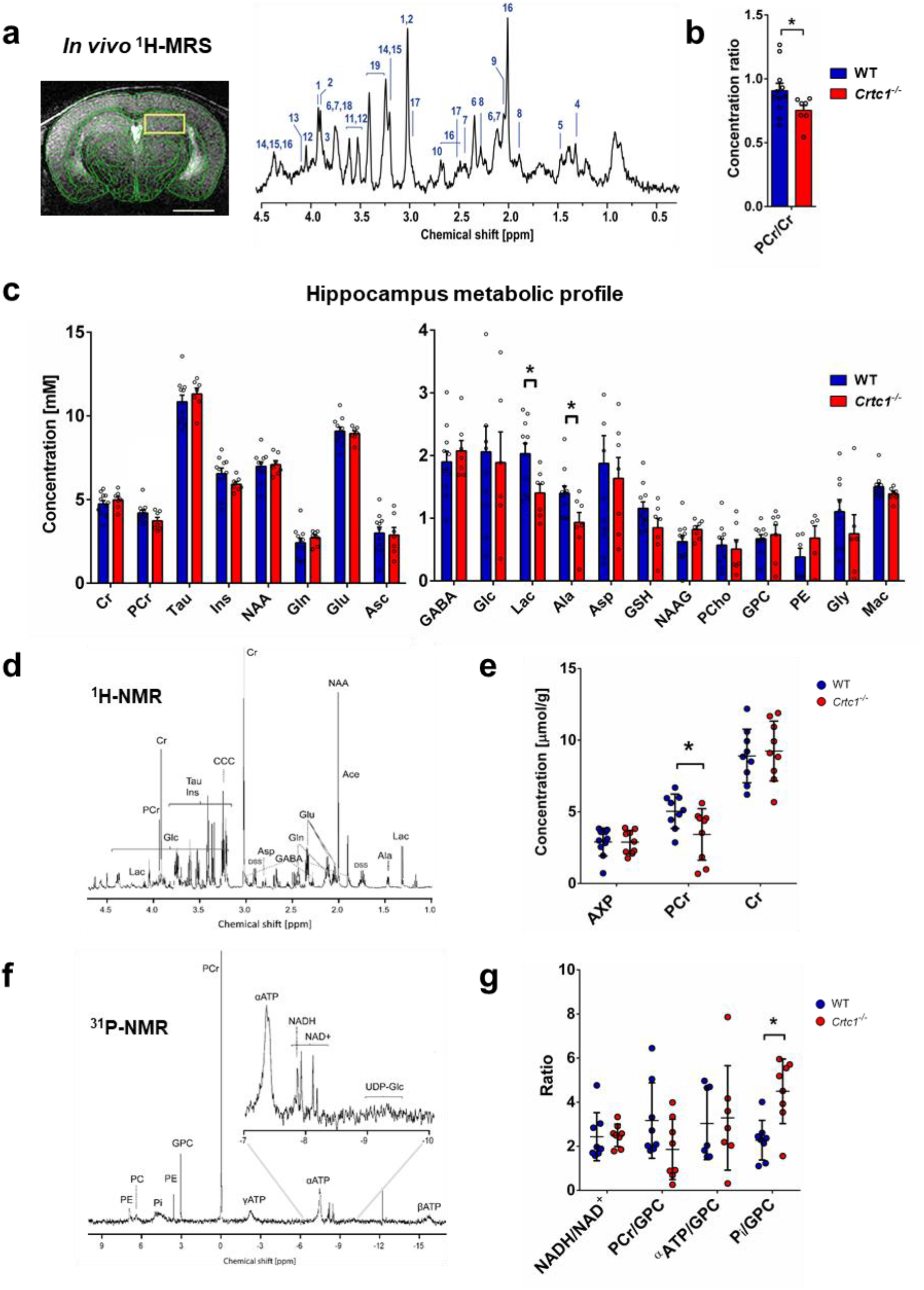
Deletion of *Crtc1* is associated with a neuroimaging fingerprint of reduced hippocampal neuroenergetics. **a**, T_2_-weighted image acquired for localized MRS (VOI including dorsal hippocampus: yellow rectangle), with a scale bar of 2mm (left) and typical ^1^H-MRS spectrum acquired in the dorsal hippocampus (DH) of 6 weeks old mice at 14.1 Tesla (right). Metabolites in the spectrum include: 1. phosphocreatine (PCr), 2. creatine (Cr), 3. glucose (Glc), 4. lactate (Lac), 5. alanine (Ala), 6. glutamate (Glu), 7. glutamine (Gln), 8. γ-aminobutyric acid (GABA), 9. N-acetylaspartyl-glutamate (NAAG), 10. aspartate (Asp), 11. glycine (Gly), 12. myo-inositol (Ins), 13. phosphoethanolamine (PE), 14. glycerophosphorylcholine (GPC), 15. phosphorylcholine (PCho), 16. N-acetyl-aspartate (NAA), 17. glutathione (GSH), 18. ascorbate (Asc), 19. taurine (Tau) as well as macromolecules (mac). Spectrum is shown with 3Hz exponential apodization. **b,c** Quantification of DH neurochemical profile from ^1^H-MRS in wild-type (WT; n=10) and *Crtc1*^-/-^ (n=6) mice, \**P*<0.05, unpaired Student’s t-test. Data are shown as mean±s.e.m. **d**, Typical high-resolution ^1^H-NMR spectrum of DH extracts acquired at 600 MHz with **e**, quantification of AXP (sum of AMP, ADP and ATP), PCr and Cr in wild-type (n=8) and *Crtc1*^-/-^ (n=8) mice, \**P*<0.05, Mann-Whitney test. **f**, Typical high-resolution ^31^P-NMR spectrum of DH extracts with **g**, quantification of NADH/NAD^+^ ratio as well as PCr, α-ATP and inorganic phosphate (Pi) relative to the GPC resonance in wild-type (n=8) and *Crtc1*^-/-^ (n=8) mice, \**P*<0.05, Mann-Whitney test. All high-resolution data (e and g) are shown as mean±s.d.

### Deletion of *Crtc1* impacts hippocampal glycolytic metabolism with subsequent mitochondrial allostatic load

We next aimed to identifying the origin of hippocampal metabolic alterations by assessing glycolytic and mitochondrial energetic function. Measuring brain glucose utilization with PET, upon infusion of ^18^F-fluorodeoxyglucose (^18^FDG) radiotracer, revealed that *Crtc1*^-/-^ mice have less glucose consumption in the hippocampus compared to controls (Fig.2a-d). Accumulation curves of ^18^F in hippocampus, resulting from cellular incorporation of ^18^FDG into ^18^FDG-6P through the action of hexokinase, were clearly reduced in the *Crtc1*^-/-^ mice (Fig.2a-b), which was associated with a 20% lower cerebral metabolic rate of glucose obtained by mathematical modeling (CMR_Glc_; Fig.2c-d; *P*=0.0045). Interestingly, the ability to produce energy through mitochondrial function did not appear to be affected *per se*, as we did not observe any significant alteration of electron transport system (ETS) expression (mtDNA- or nDNA-encoded; Fig.2e) or respiration efficiency (Fig.2f) in *Crtc1*^-/-^ mice. Furthermore, no apparent difference in master regulators of mitochondrial biogenesis and function, i.e. PGC1α and β (peroxisome proliferator-activated receptor gamma coactivator 1α and β), was observed (Fig.2g), strengthening the idea that mitochondrial capacity is not directly affected by deletion of *Crtc1*. Nevertheless, the low PCr/Pi ratio described earlier strongly suggests that mitochondria are under pressure to maintain homeostasis, as supported by creatine kinase (cytoplasmic, *Ckb*; and mitochondrial, *Ckmt1*) upregulation in *Crtc1*^-/-^ mice (Fig.2g). In sum, these results suggest that the low hippocampal PCr and Lac content observed in young *Crtc1*^-/-^ mice arises from impaired glycolytic metabolism, creating a pressure to maintain steady ATP levels (Fig.2h), a situation described as an allostatic load.

**Figure 2:**
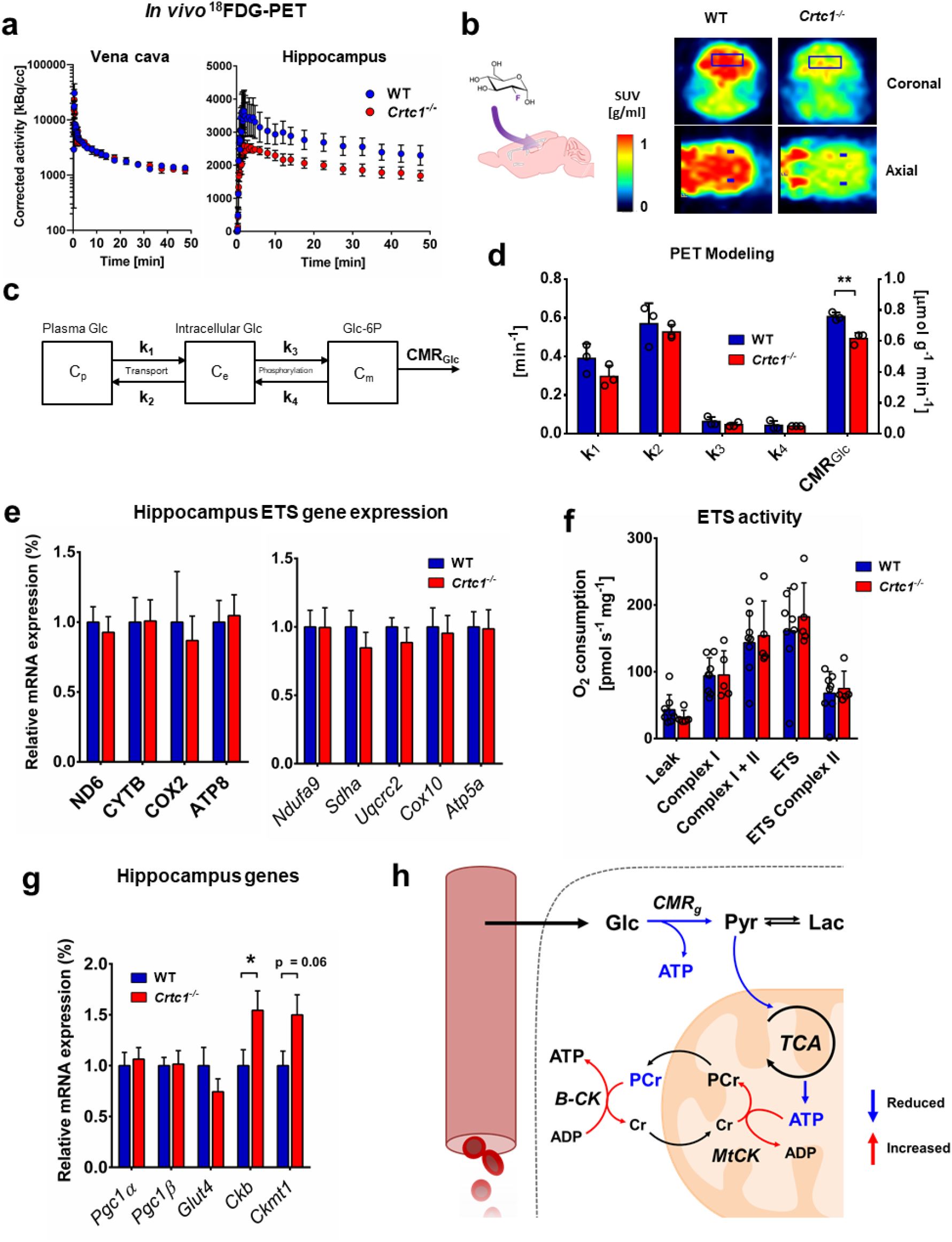
Deletion of *Crtc1* impacts hippocampal glycolytic metabolism with subsequent mitochondrial allostatic load. **a-d**, *In vivo* ^18^FDG-PET results show reduced glycolytic activity in the hippocampus of in *Crtc1*^-/-^ mice compared to wild-type (WT) mice. **a**, Time course of the radioactive decay-corrected activity, to the start of the acquisition, for vena cava (left) and hippocampus (right) in wild-type (n=3) and *Crtc1*^-/-^ (n=3) mice. **b,** Schematic of brain ^18^FDG uptake (left) and heat-maps of standard uptake values (SUVs) at steady-state (last 5min) after ^18^FDG delivery in one *Crtc1*^-/-^ and wild-type mouse (right). **c,** Mathematical model used for assessing glucose entry and metabolism from PET data. Glucose (Glc) is in exchange between one plasma (Cp) and one intracellular (Ce) pool with kinetic constants k1 and k2. A glucose-6-phosphate (Glc-6P) pool (Cm) is produced from phosphorylation of intracellular Glc via kinetic constants k3 and k4. Glc-6P is then further metabolized through glycolysis, referred to here as the “cerebral metabolic rate of glucose” (CMR_Glc_), **d**, Glucose metabolism parameter estimates from mathematical modeling of hippocampal ^18^FDG-PET data. \*\**P*<0.005, unpaired Student’s t-test. **e-g,** Mitochondrial status is not directly affected by deletion of *Crtc1*. **e**, Relative electron transfer system (ETS) gene expression in dorsal hippocampus of wild-type (n=9) and *Crtc1*^-/-^ (n=7) mice. mtDNA-encoded: ND6, complex I; CYTB, complex II; COX2, complex IV; ATP8, complex V. nDNA-encoded: *Ndufa9*, complex I; *Sdha*, complex II; *Uqcrc2*, complex III; *Cox10*, complex IV; *Atp5a*, complex V. **f**, Mitochondrial respirometry in dorsal hippocampus of wild-type (n=8) and *Crtc1*^-/-^ (n=5) mice. **g**, Mitochondrial gene expression in dorsal hippocampus of wild-type (n=9) and *Crtc1*^-/-^ (n=8) mice. *Pgc1α* and *β,* Peroxisome proliferator-activated receptor gamma coactivator 1 alpha and beta; *Ckb*, creatine kinase B-type; *Ckmt1*, creatine kinase mitochondrial type. **h**, Schematic representation of hippocampal mitochondrial allostatic load. Reduced glycolytic function leads to fewer pyruvate available for oxidation in the mitochondria. The resulting lack of ATP produced from mitochondria and glycolysis is compensated by higher PCr hydrolysis, which helps buffer ATP depletion to maintain homeostasis and potentially stimulated by the upregulation of creatine kinases expression. Glc, glucose; Pyr, pyruvate; B-CK, cytoplasmic creatine kinase; MtCK, mitochondrial creatine kinase. All data are shown as mean±s.e.m.

### Hippocampal energetic status reflects the depressive-like behavior of *Crtc1*^-/-^ mice

To test the stability over time of these hippocampal energetic alterations and determine if they were associated with the depressive-like behavior of *Crtc1*^-/-^ mice, we subjected WT and *Crtc1*^-/-^ animals to social isolation from the age of 6 weeks and monitored their neurochemical profile and behavior longitudinally (Fig.3a). Social isolation was used to ensure a comparable social environment between groups and reduce aggression-related effects within cages^28^. A higher level of depressive-like behavior was observed (Fig.3b) for *Crtc1*^-/-^ mice under basal conditions (6 weeks of age) as reflected in forced swim test (FST; *P*=0.02) but not in tail suspension test (TST; n.s). Surprisingly, 18 weeks of social isolation had an opposite effect on the behavior of the two groups (averaged z-scores, Interaction: F_2,28_=10.26, *P*=0.0005; TST, Interaction: F_2,28_=5.16, *P*=0.012; FST, Interaction: F_2,28_=3.87, *P*=0.035; two-way ANOVA). Moreover, an inversion in the hippocampal energetic profile (Fig.3c) coincided with this switch in behavior (Lac, Interaction: F_2,28_=7.32, *P*=0.003; PCr, Interaction: F_2,28_=4.78, *P*=0.017; PCr/Cr, Interaction: F_2,28_=2.79, *P*=0.08; two-way ANOVA). Interestingly, hippocampal glucose concentration rose only in *Crtc1*^-/-^ mice upon social isolation (Time effect: F_2,28_=3.43, *P*=0.050, two-way ANOVA; *Crtc1*^-/-^ 6-weeks vs. 6-months, \**P*<0.05, Bonferroni’s test). We then performed correlational analyses to further relate metabolite hippocampal markers with behavior (Fig.3d) and found a significant negative correlation between the depressive-like behavior and Lac (Lac vs. averaged z-scores: R=-0.35, p=0.01). To test whether these metabolic modifications were associated with a change in gene expression, we analyzed relative mRNA content in DH at the end of the protocol (Fig.3e and Fig.S2a) and found a difference in *Pgc1α* (*P*=0.04) and *Glut4* (*P*=0.01) between the two groups, while creatine kinases levels were no longer significantly different (n.s.). Notably, differences in depressive-like behavior between *Crtc1*^-/-^ and wild-type mice were not related to locomotor activity at any age (Fig.S2b, n.s.) or PFC volume and tCho content, which both correlated with each other and remained increased in *Crtc1*^-/-^ independently of the animal’s age (Fig.S2c and Fig.S3; tCho, Genotype effect: F_1,148_=12.89, *P*=0.003; Volume, Genotype effect: F_1,42_=14.61, *P*=0.0004; two-way ANOVA; Correlation: R=0.31, p=0.03). Importantly, social isolation stimulated the development of a MeS-related phenotype in both groups as suggested by the rise in body weight (Fig.3f), which developed faster over time for *Crtc1*^-/-^ mice (Genotype effect: F_1,14_=5.84, *P*=0.01; Interaction: F_2,28_=5.11, *P*=0.03, two-way ANOVA), and the high level of blood MeS markers (insulin, glucose and triglyceride), which were not significantly different between the groups (Fig.3g; n.s.). Overall, these results confirm that the hippocampal neuroenergetic status of *Crtc1*^-/-^ mice reflects their depressive-like behavior and indicate an apparent dependence on the experienced environment.

**Figure 3:**
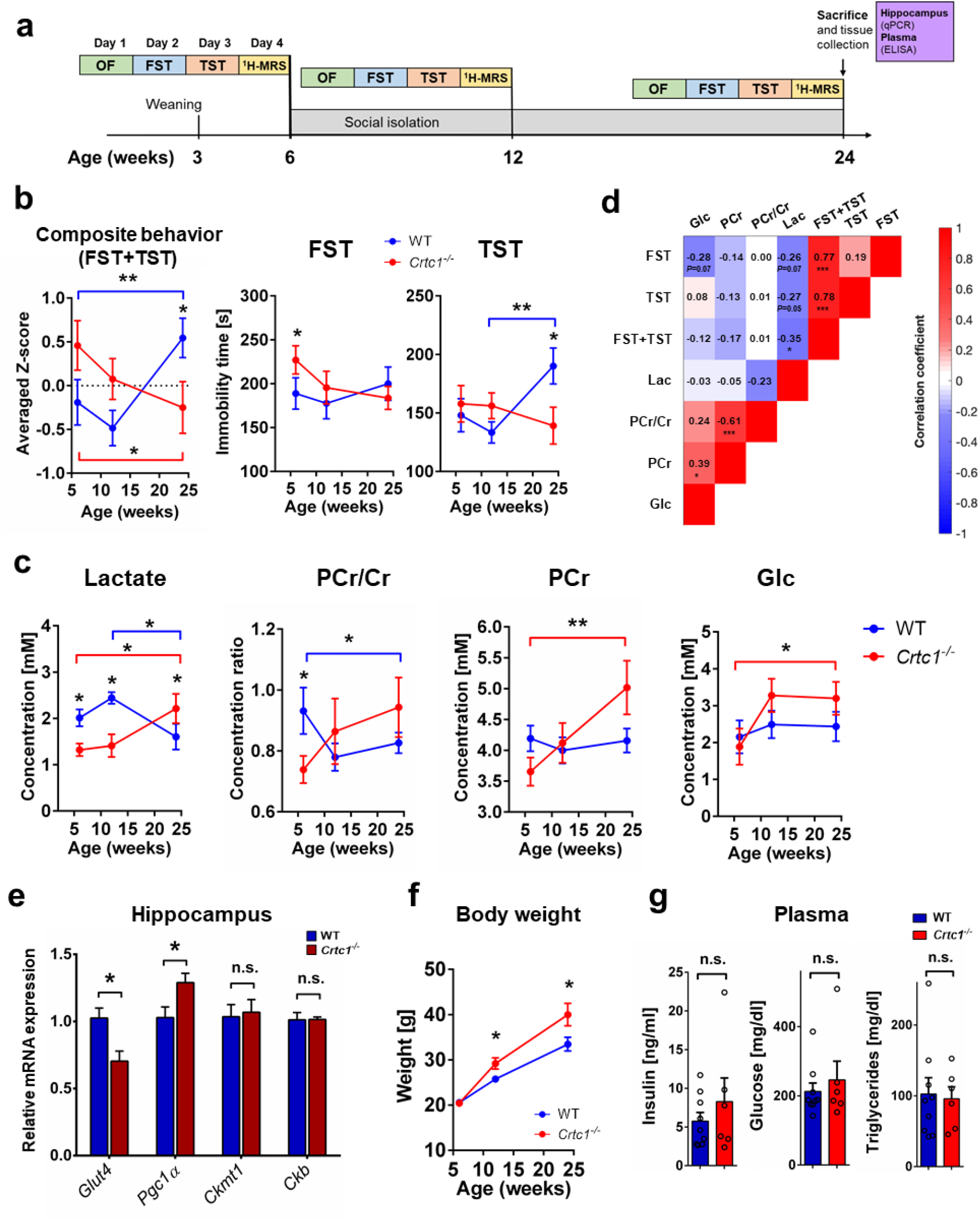
Hippocampal energetic status reflects the depressive-like behavior of *Crtc1*^-/-^ mice. **a**, Experimental design, and timeline of the longitudinal protocol used involving social isolation. wild-type (WT; n=10) and *Crtc1*^-/-^ (n=6) mice underwent a set of behavioral tests including an open-space test (OF; day 1), a forced swim test (FST; day 2) and a tail-suspension test (TST; day 3) followed by a ^1^H-MRS scan on day 4. After this first set of experiments, animals were isolated at the age of 6 weeks and the whole procedure was repeated at 12 and 24 weeks of age. After the last ^1^H-MRS scan, animals were sacrificed, and hippocampal and plasma samples were collected for analysis. **b**, A switch in depressive-like behavior between *Crtc1*^-/-^ and wild-type mice occurs after 18 weeks of social isolation as revealed by the inversion in immobility time in TST (right panel; Interaction: F_2,28_=5.16, *P*=0.012), FST (center panel; Interaction: F_2,28_=3.87, *P*=0.035) and averaged z-score of TST and FST (left panel; Interaction: F_2,28_=10.26, *P*=0.0005). Two-way ANOVA, followed by Fisher LSD posthoc test; \**P*<0.05, \*\**P*<0.01. **c**, Hippocampal neuroenergetic profile switches between *Crtc1*^-/-^ and wild-type mice at the end of 18 weeks of isolation as revealed by the inversion of lactate concentration (left panel; Interaction: F_2,28_=7.32, *P*=0.003), PCr/Cr ratio (center left panel; Interaction: F_2,28_=2.79, *P*=0.08) and PCr (center right panel; Interaction: F_2,28_=4.78, *P*=0.017;). Hippocampal glucose levels increased in the *Crtc1*^-/-^ group only at the end of the 18 weeks of isolation (Time effect: F_2,28_=3.43, *P*=0.050). Two-way ANOVA, followed by Bonferroni’s posthoc test; \**P*<0.05, \*\**P*<0.01. **d**, Correlative analysis between depressive-like behavior and hippocampal energy metabolite content. A significant negative correlation between Lac and behavior was found when results from FST and TST were considered together (R=-0.351, p=0.013). Color code represents Pearson’s correlation coefficient and the analysis included all longitudinal age time points. Pearson’s Rs are shown for each correlation with associated *P* value (uncorrected for multiple comparisons); *\*P*<0.05, *\*\*\*P*<0.0001. **e**, At the end of 18 weeks of isolation, hippocampal levels of *Pgc1α* mRNA were higher while *Glut4* levels were lower in *Crtc1*^-/-^ as compared to wild-type mice. Mitochondrial and cytoplasmic creatine kinases were not significantly different (n.s.) between the two groups. Unpaired Student’s t-test, \**P*<0.05. **f**, Body weight of all animals increased significantly over time (Time effect: F_2,28_=123.2, *P*<0.0001) but increased more in the *Crtc1*^-/-^ group (Genotype effect: F1,14=5.84, *P*=0.030; Interaction: F_2,28_=5.11, *P*=0.013). Two-way ANOVA followed by Fisher LSD post-hoc test \**P*<0.05. **g**, Plasma markers of metabolic syndrome (insulin, glucose and triglycerides) were high in both groups but not significantly different from each other (n.s.).

### Restoring hippocampal energy balance with energy-boosting ebselen mood-stabilizer rescues depressive-like behavior in *Crtc1*^-/-^ mice

Social isolation appeared to be beneficial for *Crtc1*^-/-^ mice, consistent with their known aggressive behavior and social impairments towards other individuals^26^. We thus hypothesized that a repeated open-space forced swim test (OSFST) protocol (Fig.4a), which contains an environmental-rather than social-stress component, would challenge neuroenergetics in both groups of mice. In parallel, we tested whether improving brain metabolism with an energy-stimulating compound would reverse the stressful effects of the OSFST. To maximize the translational relevance of our findings, we decided to treat animals by oral administration of ebselen, a neuroprotective and antioxidant compound^29^ with comparable pharmacological properties as lithium (i.e. inhibitor of GSK3β and inositol monophosphatase (IMP))^30^ and with a strong clinical potential^31,32^. After 4-consecutive days of swimming sessions and establishment of a stable depressive-like behavior in all groups of mice, animals were treated with either ebselen or vehicle twice a day for 3 weeks. As expected, the depressive-like behavior, measured as immobility time in OSFST, was higher in *Crtc1*^-/-^ mice over time (Fig.4b; Genotype effect: F_1,10_=65.09, *P*<0.0001, two-way ANOVA). Ebselen rescued the behavior of *Crtc1*^-/-^ mice (Interaction: F_1,10_=41.84, *P*<0.0001; Treatment effect: F_1,10_=5.45, *P*=0.04, two-way ANOVA) and led to an improvement in hippocampal energy metabolism (Fig.4c,d). More specifically, ebselen raised hippocampal PCr content (Fig.4d; ΔPCr/Cr, Treatment effect: F_1,31_=4.41, *P*=0.04; two-way ANOVA) compared to the untreated groups, but lowered lactate levels in *Crtc1*^-/-^ mice at the end of the study protocol (Fig.4c; Lac day 21, *P*=0.045; unpaired t-test), in line with enhanced mitochondrial activity. Furthermore, the difference in energy metabolite content correlated with a difference in behavior (ΔPCr/Cr, R=-0.54, *P*=0.02; ΔLac, R=0.41, *P*=0.01) suggesting that both events were linked (Fig.4e). Gene expression analysis (Fig.4f and Fig.S4a) supports that ebselen stimulated DH mitochondrial function through inhibition of GSK3β, as highlighted by a treatment effect observed in *Pgc1α* (F_1,27_=13.28, *P*=0.009), *Glut4* (F_1,27_=8.22, *P*=0.001) and *Ckmt1* (F_1,27_=4.79, *P*=0.04, two-way ANOVA) mRNA content. Importantly, ebselen did not interfere with the increased body weight and high insulin and triglyceride levels in *Crtc1*^-/-^ mice (Fig.4g; Treatment effect, n.s), confirming a brain-specific mechanism, neither did ebselen affect PFC volume and tCho concentration differences observed in *Crtc1*^-/-^ mice (Fig.S4b-d). Finally, to assess the potential clinical relevance of the identified neuroimaging markers we determined their specificities and sensitivities using receiver operating characteristic (ROC) curves (Fig.S4e-f). Prefrontal volume and tCho concentration were able to differentiate *Crtc1*^-/-^ mice from their wild-type counterparts with an area under the curve (AUC) of up to 82% (95% CI 0.755-0.886), when combined into an averaged z-score. The ability of hippocampal neuroenergetic markers to differentiate mice with ‘high’ levels of depressive-like behavior from those with ‘low’ levels was more modest, with an AUC of up to 66% (95% CI 0.555-0.756), when combined into an averaged z-score. In summary, stimulating mitochondrial energy metabolism was able to rescue the depressive-like behavior induced by stress in *Crtc1*^-/-^ mice, leading to neuroimaging-based modifications that followed the treatment response.

**Figure 4:**
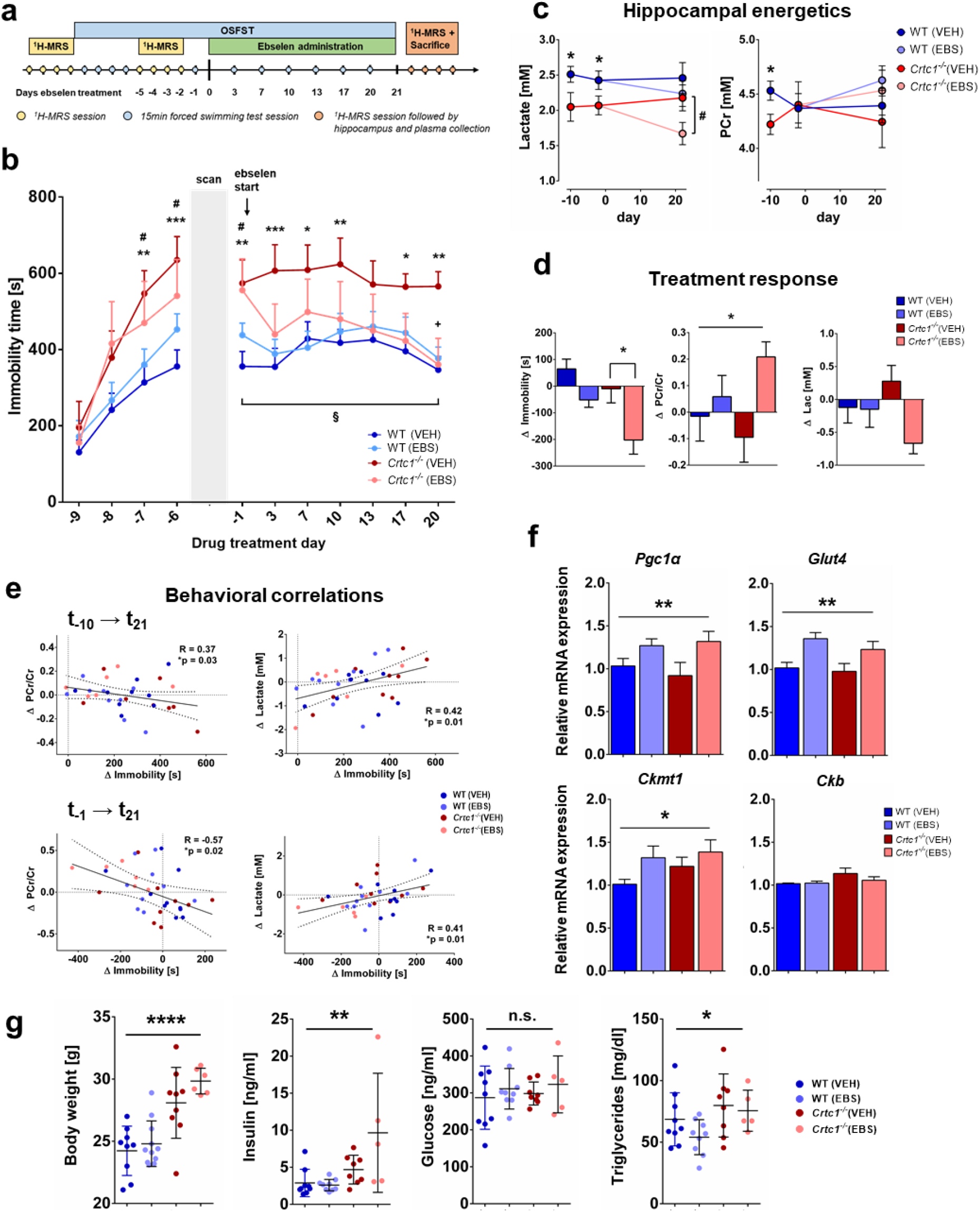
Restoring hippocampal energy balance with energy-boosting ebselen mood-stabilizer rescues depressive-like behavior in *Crtc1*^-/-^ mice. **a**, Experimental design, and timeline of the ebselen treatment protocol during open-space forced swim test (OSFST). First, animals underwent a single basal ^1^H-MRS scan (during days -13 to -10), followed by 4 consecutive forced swimming sessions (day -9 to -6). During days -5 to -2, animals underwent a second ^1^H-MRS scan and a fifth swimming session on day -1, prior to the treatment start. Animals were administered ebselen (wild-type(EBS), n=8; *Crtc1*^-/-^(EBS), n=6) or vehicle (wild-type(VEH), n=9; *Crtc1*^-/-^ (VEH), n=9) twice daily from day 0 and until the end of the OSFST protocol (day 21), while swimming sessions were repeated regularly every 3-4 days. A final ^1^H-MRS scan was performed at the end of the study (between days 22 to 25), with subsequent hippocampal and plasma collection for analyses. **b**, Depressive-like behavior in the OSFST was higher in the *Crtc1*^-/-^ mice (Genotype effect: F1,10=65.09, *P*<0.0001) but reduced by ebselen (Treatment effect: F1,10=5.45, *P*=0.04, interaction: F1,10=41.84, *P*<0.0001). Immobility of all *Crtc1*^-/-^ mice was increased after the first 4 days swimming session (\*\**P*<0.01, \*\*\**P*<0.005 for VEH and ^#^*P*<0.05 for EBS vs their respective wild-type group). Depressive-like behavior of *Crtc1*^-/-^ VEH group remained significantly higher than wild-type over the 21 days of test (\**P*<0.05, \*\**P*<0.01 and \*\*\**P*<0.005 for VEH *Crtc1*^-/-^ vs VEH wild-type). After the 21 days of OSFST, the depressive-like behavior of the treated *Crtc1*^-/-^ animals was significantly reduced (^+^*P*<0.05 compared to *Crtc1*^-/-^ VEH at day 20 and ^§^*P*<0.05 compared to *Crtc1*^-/-^ EBS at day 0). Two-way ANOVA for repeated measures, followed by Fisher LSD posthoc test. **c**, Hippocampal energy metabolite concentrations during the OSFST protocol. Lactate and PCr content were lower in *Crtc1*^-/-^ animals relative to wild-type animals under basal conditions (days -13 to -10; Unpaired Student’s t-test, \**P*<0.05; wild-type, n=22; *Crtc1*^-/-^, n=16), but only lactate remained lower in *Crtc1*^-/-^ animals after ebselen treatment (days 22 to 25; Unpaired Student’s t-test, ^#^*P*<0.05; wild-type(EBS), n=8; *Crtc1*^-/-^(EBS), n=6). **d**, Ebselen treatment reduced depressive-like behavior and increased hippocampal high-energy phosphate content (difference between day 21 and day 0). Ebselen treatment increased the PCr/Cr ratio (Treatment effect: F1,31=4.41, \**P*=0.044, two-way ANOVA) and tended to reduce lactate (Treatment effect: F1,31=3.49, *P*=0.071, two-way ANOVA), together with immobility reduction (Treatment effect: F1,31=13.7, *P*=0.0008; Genotype effect: F1,31=7.30, *P*=0.011; two-way ANOVA, followed by Bonferroni’s test, \**P*<0.05). **e**, (top) The increase in immobility from baseline (day -10) as a result of OSFST (day 21) correlated with a reduction in PCr/Cr (R=-0.37, \**P*=0.03) and rise in lactate (R=0.42, \**P*=0.01). (bottom) Immobility reduction (from day -1) as a result of treatment (day 21) correlated with a rise in PCr/Cr (R=-0.57, \**P*=0.02) and a drop of lactate (R=0.41, \**P*=0.01). **f**, Ebselen induced expression of energy-related genes in hippocampus. Treated animals had higher mRNA level of *Pgc1α* (Treatment effect: F_1,27_=13.28, \*\**P*=0.0087), *Glut4* (F_1,27_=8.22, \*\**P*=0.0011) and *Ckmt1* (F_1,27_=4.79, \**P*=0.037). Relative cytoplasmic creatine kinase (*Ckb*) expression tend to increase in *Crtc1*^-/-^ mice at the end of the OSFST protocol (Genotype effect: *Ckb*: F_1,27_=3.78, *P*=0.06; *Ckmt1*: F_1,27_=1.59, *P*=0.21). Two-way ANOVA. **g**, *Crtc1^-/-^* showed significantly higher body weight (Genotype effect: F1,31=37.7, \*\*\*\**P*<0.0001), plasma insulin (F_1,27_=12.24, \*\**P*<0.0016) and triglyceride levels (F_1,27_=4.78, \**P*=0.038) at the end of the OSFST protocol, with not treatment effect (n.s.). Two-way ANOVA; n.s., not significant. Data are reported as mean±s.e.m.

### GABAergic dysfunction links impaired hippocampal glucose metabolism with depressive-like behavior in *Crtc1*^-/-^ susceptible mice

Finally, to assess the relative brain cellular metabolic contributions, we acquired indirect ^13^C-carbon magnetic resonance spectroscopy (^1^H-[^13^C]-MRS; Fig.5a) data to assess metabolic fluxes using mathematical modeling. Fractional isotopic ^13^C-enrichment (FE) of brain glucose and downstream metabolites revealed clear group differences in animals of 6 weeks in age (Fig.5b) involving metabolites associated with glycolysis (U-Glc, LacC3), tricarboxylic acid (TCA) cycle (GluC4) and GABAergic neurons metabolism (GABAC2-4). When fitting the mathematical models to the ^13^C-labeling data (Fig.5c and Fig.S5a), we found that reduced glucose consumption (i.e. CMR_Glc_) led to a drop of TCA cycle activity in both excitatory (−36%, *P*<0.0001) and inhibitory (−14%, *P*=0.01) neurons of *Crtc1*^-/-^ mice. Neurotransmission flux was overall increased (Fig.S5b; V_NT_=0.06±0.01 for wild-type vs. 0.09±0.02 μmol/g/min for *Crtc1^-/-^, P*=0.004) when considering metabolism as a whole (1-compartment) but analysis with a more complex model (pseudo 3-compartment model, i.e. that considers the relative cellular metabolic contributions) indicated this effect was more pronounced in GABAergic (V_NT_^i^+V_ex_^i^; 6-fold increase) than glutamatergic (V_NT_^e^; 2-fold increase) neurotransmission. Importantly, the increase in GABA labeling (Fig.6b) did not arise from an increase in GAD activity according to our model (V_GAD_=0.32±0.06 for wild-type vs. 0.30±0.08 μmol/g/min for *Crtc1*^-/-^, n.s.) but reflected a dilution originating from exchange between two GABA pools and possibly triggered by GABAergic neurotransmission recycling (Fig.S5c; V_ex_^i^=0.0006±0.0002 for wild-type vs. 0.007±0.003 μmol/g/min for *Crtc1^-/-^, P*=0.02), in line with a probable inhibitory neuron hyperactivity. Furthermore, despite the relatively higher drop of ATP production rate in excitatory (−35%) compared to inhibitory (−15%) neurons (Fig.S5d), the relative oxidative allostatic load calculated as the neurotransmission relative to ATP production (see methods) indicated a ~2.7-fold higher load for inhibitory neurons (8.4x higher in *Crtc1*^-/-^ mice) relative to excitatory neurons (3.1x higher in *Crtc1*^-/-^ mice), suggesting that GABAergic inhibitory neurons might be more at risk.

**Figure 5:**
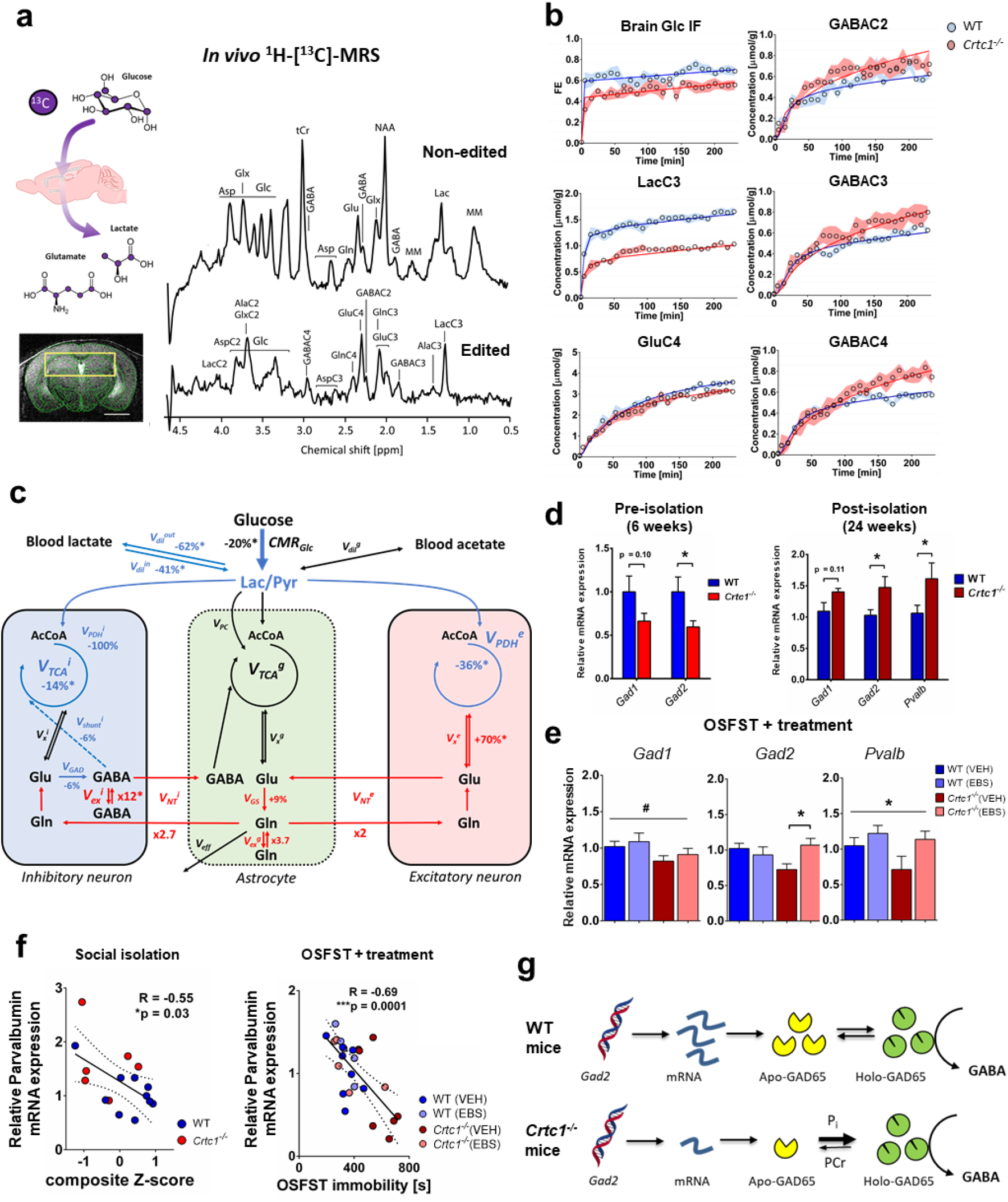
GABAergic dysfunction links impaired hippocampal glucose metabolism with depressive-like behavior in *Crtc1*^-/-^ susceptible mice. **a**, Schematic of ^13^C-labelled glucose brain uptake and subsequent metabolite labeling (upper left) ^1^H-[^13^C]-MRS spectra acquired in the bilateral dorsal hippocampus of a 6 weeks old WT mouse (right) as shown with the selected VOI (yellow box) on the associated MRI image (lower left). The non-edited spectrum (top) shows the total metabolic profile, while the edited spectrum (bottom) identifies the fraction of metabolites that have incorporated ^13^C-labeling. Scale bar=2mm. **b**, Fractional isotopic ^13^C-enrichment (FE) of glucose and key metabolites in the hippocampus during ^1^H-[^13^C]-MRS experiment. Fitting of the data with a pseudo 3-compartment model of brain glucose metabolism is shown with a straight line for wild-type (WT; in blue) and *Crtc1*^-/-^ (in red) mice. 6 weeks old wild-type (n=8) and *Crtc1*^-/-^(n=8). Data presented as mean±s.d. **c**, Schematic representation of hippocampal glucose utilization differences between wild-type and *Crtc1*^-/-^ mice after metabolic flux analysis using a pseudo 3-compartment model. Metabolic fluxes that were higher in *Crtc1*^-/-^ animals (compared to their wild-type littermates) are shown in red, while those found lower are shown in blue and those found without any difference or fixed during the modeling remain in black. Cerebral metabolic rate of glucose (CMR_Glc_); brain lactate influx (V_dil_^in^) and outflux (V_dil_^out^) from blood; pyruvate dilution flux (V_dil_^g^); excitatory neuron TCA cycle (V_PDH_^e^); inhibitory neuron pyruvate dehydrogenase activity (V_PDH_^i^); GABA shunt flux (V_shunt_^i^); inhibitory neuron TCA cycle (V_TCA_^i^=V_PDH_^i^+V_shunt_^i^); glial pyruvate carboxylase (V_PC_); glial TCA cycle (V_TCA_^g^); excitatory neuron (V_x_^e^), inhibitory neuron (V_x_^i^) and glial (V_x_^g^) transmitochondrial fluxes; excitatory neurotransmission flux (V_NT_^e^); inhibitory neurotransmission flux (V_NT_^i^); glutamate decarboxylase activity (VGAD); Gln exchange flux (V_ex_^g^); GABAergic exchange flux (V_ex_^i^); glutamine synthetase activity (V_GS_) and Gln efflux (V_eff_). Relative flux increase/decrease is indicated for *Crtc1^-/-^* mice compared to WT littermates, as calculated from fluxes in μmol/g/min from Fig.S5c; and an asterisk (*) indicates a statistically significant difference between the two groups. . **d**, GABAergic gene expression (*Gad1, Gad2* and parvalbumin (*Pvalb*)) in the hippocampus under basal conditions (6 weeks age; left) or after social isolation (24 weeks age; right). Unpaired Student t-test, \**P*<0.05; basal, wild-type (n=6) and *Crtc1*^-/-^ (n=6); longitudinal, wild-type (n=10) and *Crtc1*^-/-^ (n=6). **e**, Hippocampal gene expression of *Gad1, Gad2* and *Pvalb* after the OSFST protocol (wild-type(VEH), n=9; wild-type(EBS), n=8; *Crtc1*^-/-^(VEH), n=9; *Crtc1*^-/-^(EBS), n=6). *Gad1* was significantly reduced in the *Crtc1*^-/-^ group (Genotype effect: F_1,28_=4.39, \**P*=0.045, two-way ANOVA), while ebselen treatment increased the levels of *Gad2* (Interaction: F_1,27_=5.53, \**P*=0.026, two-way ANOVA; \**P*<0.05, Bonferroni’s post-hoc test) and parvalbumin (Treatment effect: F_1,24_=4.28, \**P*=0.049, two-way ANOVA). **f**, Correlation between depressive-like behavior and level of *Pvalb* expression in the hippocampus after social isolation (left; 24 weeks of age; R=-0.55, *P*=0.03) and OSFST protocols (right; 10 weeks of age; R=-0.69, *P*=0.0001). The dotted lines represent the 95% confidence interval of the linear regression line. **g**, Scheme of potential relation between GAD expression level, energy metabolite binding and enzyme activity.

**Figure 6:**
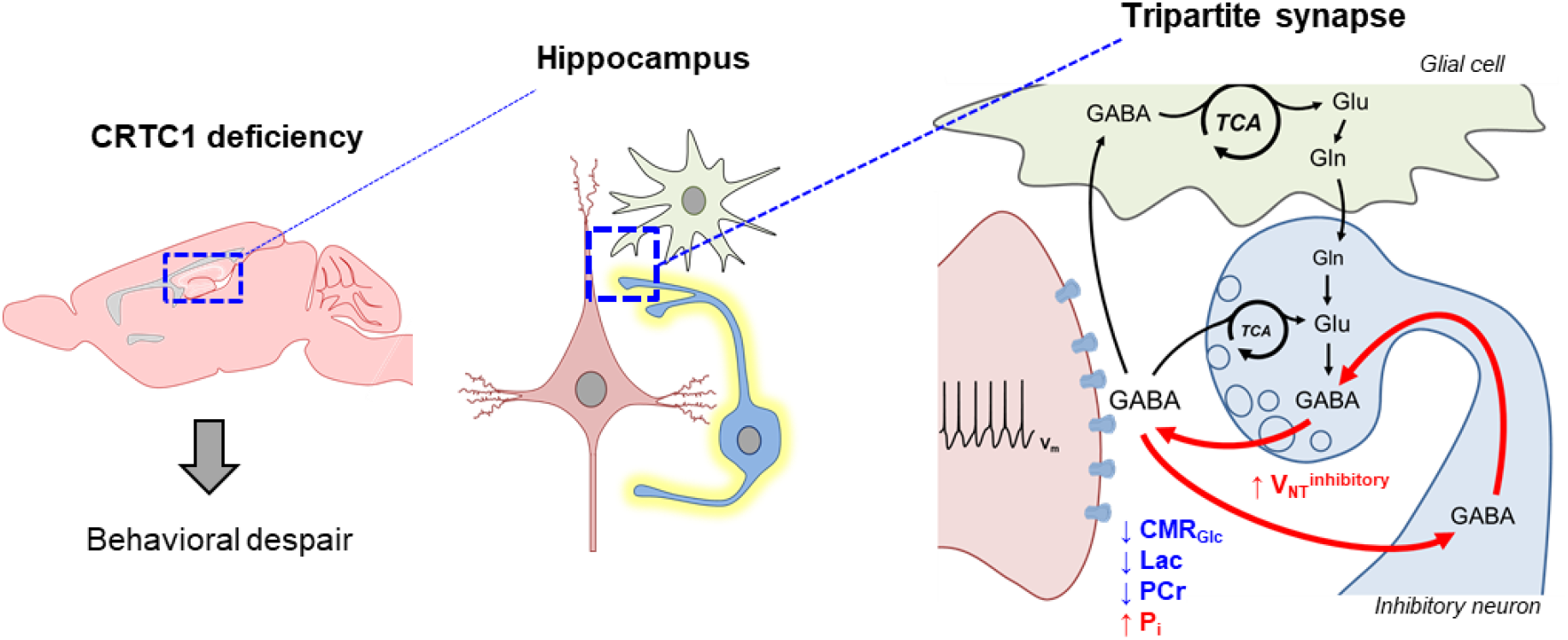
Scheme of hippocampal GABAergic hyperactivity resulting from low energetic status linking *Crtc1* deletion to depressive-like behavior. Reduced hippocampal glucose metabolism capacity relative to neuronal neurotransmitter cycling-demands leads to low energetic status (high inorganic phosphate (Pi) and low phosphocreatine (PCr) levels) in *Crtc1* deficient mice. This results in excessive GABAergic neurotransmitter cycling and depressive-like behavior.

To further determine if the GABAergic system is particularly impacted by hippocampal energetic impairments, we re-analyzed main GABAergic gene expression in our different experimental protocols. Interestingly, levels of *Gad2* and parvalbumin (*Pvalb*) were strongly associated with the behavioral state of the animals (Fig.5d-f). At the age of 6 weeks, *Gad2* was lower in *Crtc1*^-/-^ mice (*P*=0.04) when depressive-like behavior was high (Fig.3b), while it was increased after social isolation (*P*=0.03; Fig5.d) when the behavior was inverted as well (Fig.3b). Importantly, ebselen restored the levels of both *Gad2* (Interaction: F_1,27_=5.53, *P*=0.03) and *Pvalb* (treatment effect: F_1,24_=4.28, p=0.049) in *Crtc1*^-/-^ mice after OSFST. Finally, *Pvalb* was the only gene that correlated directly with the level of depressive-like behavior in both experiments (Social isolation: R=-0.55, *P*=0.03; OSFST+treatment: R=-0.69, *P*=0.0001). The above results suggest that the hippocampal GABAergic system might be mechanistically involved in the depressive-like behavior induced by neuroenergetic impairments.

## Discussion

Understanding how genetic and environmental factors interact in metabolic diseases and how they impact normal brain and behavior is central for better diagnosing and treating related mood disorders. Because of its central role in regulating brain metabolism and its strong association with features of MeS in psychiatric patients^11–14^, *Crtc1* is a key candidate gene to understand how (neuro-)metabolic alterations can affect normal behavior. In this study, we have been able to identify reduced hippocampal energy metabolism in *Crtc1*-deficient mice that translated into measurable *in vivo* neuroimaging markers. We have demonstrated that these neurochemical impairments were associated with animal depressive-like behavior, which could be reversed with an energy-boosting treatment known for its mood-stabilizing properties. Finally, we provide evidence for a hyper-activation and allostatic load of the hippocampal GABAergic system that could mediate behavioral consequences of the observed neuroenergetic imbalance.

Even though *Crtc1* is predominantly expressed in the brain^26,33^ deleting this gene in mice induces a systemic metabolic deregulation^22^, such as insulin resistance and obesity, together with a depressive-like phenotype^26,27^. While the association of MeS and behavioral alterations is likely to be complex and multifactorial, we report a clear link between brain glucose uptake and depressive-like behavior. Specifically, low glycolytic activity in *Crtc1*^-/-^ mice was associated with reduced levels of lactate and increase in high-energy phosphate hydrolysis (i.e. high level of Pi and low levels of PCr) in hippocampus that correlated well with animal behavior (Fig.1). Importantly, our results indicate reduced hippocampal glucose uptake capacity rather than lower demand, as the neuronal activity relative to energy production (V_NT_/V_TCA_) was found to be ~3-fold higher for the *Crtc1*^-/-^ mice (Fig.5a-c and Fig.S5), pointing towards a difficulty in matching energy production with neuronal needs, or what is defined as an allostatic load^34^. This fits well with the idea that CRTC1 is required for adapting energy homeostasis according to neuronal requirements^22,35^ and is in line with several studies demonstrating the central role of brain energy metabolism in the resilience mechanisms against depressive-like behavior^36–40^. Of note, our results indicate that both glycolytic and mitochondrial pathways are fundamental for brain metabolic resilience and behavior rather than one route preferentially, as CMR_Glc_ deficiency in *Crtc1*^-/-^ mice could be compensated by an increase in oxidative metabolism. Considering that glucose entry in the brain is regulated by factors such as the insulin or IGF-1 receptors, known to influence mouse depressive-like behavior^41^, we hypothesize that reduced hippocampal glucose uptake arises from the known insulin resistance phenotype of *Crtc1*^-/-^ mice^22^. While future research will determine the exact molecular mechanisms relating *Crtc1* with brain energy capacity, our experimental data point towards the Akt/GSK3β pathway as a key player in this process. In fact, inhibition of GSK3β with ebselen improved DH energetic status and behavior through enhanced hippocampal *Pgc1α* and *Glut4* expression, with only little effect on peripheral energy markers (Fig.4g). PGC1α, as a master mitochondrial biogenesis regulator, can be inhibited through phosphorylation by GSK3β^42^, possibly impacting *Glut4* expression over the MEF2C transcription factor^43^. PGC1α has been linked with depression^44^ and bipolar disorders^45^, and its target, PPARγ, provides a plausible link between MeS and behavior. For instance, PPARγ agonists, which are well known insulin sensitizing agents^9^, show anti-depressant properties in animal models^46^ and patients^47,48^ leading to improved glucose metabolism^49^. Although studies focusing on muscle cells showed that *Crtc2*, a peripheral homolog of *Crtc1*, can induce *Pgc1α* expression^50^, we did not observe reduced *Pgc1α* levels as a result of *Crtc1* deletion (Fig.2f, Fig.3e and Fig.4f). Nevertheless, enhancing *Pgc1α* expression restored energy metabolism and behavior in *Crtc1*^-/-^ mice (Fig.4f), suggesting that CRTC1 deficiency can be compensated through different, though converging, pathways.

How then did the enhanced hippocampal energetic capacity, illustrated by higher *Pgc1α* and *Glut4* expression (Fig.4f), not affect the behavior of wild-type mice, as would be expected from this model? It is plausible that efficient energy metabolism is necessary for resilience to depressive-like behavior but is not sufficient to modulate it. Energy metabolism, either mitochondrial or glycolytic, has been widely implicated in the pathophysiological mechanisms leading to depressive-like behavior in preclinical models^36,38–40,51^ and in clinical studies^52–55^. Nevertheless, it remains unclear how altered brain energy production rates could translate into behavioral dysfunction. While several processes have been brought forward, such as metabotropic-, neuroendocrine-, inflammatory-, transcriptional-, or other responses^56–58^, our results highlight the hippocampal GABAergic neurotransmitter system as a new key player in the process linking cellular allostatic load with affected neuronal output. In fact, our metabolic flux and GABAergic gene expression analyses indicate that the inhibitory system is particularly affected by low energy status and could relate to depressive-like behavior more tightly than the level of metabolism-enhancing genes or high-energy phosphates. We have previously reported that inhibitory neurotransmission in the hippocampus has high mitochondrial oxidative dependence compared to excitatory neurotransmission in mice^59^. Accordingly, here we found that low energy production capacity in *Crtc1*^-/-^ mice was associated with a ~6-fold increase in hippocampal GABAergic neurotransmission cycling (Fig.5a-c), leading to an overall higher (~2.6-fold) oxidative allostatic load in inhibitory compared to excitatory neurons. Others have shown that GABA neuronal metabolism is highly controlled by the cellular energetic status, through the action of both GAD isoforms (GAD65 and GAD67), switching from an Apo (inactive) to a Holo (active) conformation in response to low energy metabolite concentration, i.e. increased Pi or reduced PCr or ATP^60–62^. This feature would provide a protective network-inhibition mechanism when energy demands exceed metabolic capacities. Furthermore, our present work also shows that GABAergic markers (*Gad1, Gad2* and *Pvalb*) were highly correlated with the animals’ behavior (Fig.5d-f). Considering that fast-spiking parvalbumin-positive interneurons, particularly activated during gamma-oscillations in the hippocampus, are known to be very energy consuming and mitochondria-rich^63^, improving energy metabolism might confer significant resilience to this cell population in particular. Of note, Uchida et al. reported that disruption of *Gad1* function can lead to the loss of parvalbumin neurons in the hippocampus as a result of stress exposure^64^. Interestingly, several studies have also reported lower post-mortem levels of *Gad1* expression in PFC of bipolar and schizophrenic patients^65–69^. Importantly, and of potential therapeutic relevance, ebselen was able to rescue the behavior in *Crtc1*^-/-^ mice, by restoring hippocampal energy metabolites and levels of *Gad2* and *Pvalb* expression (Fig.4). This resonates with previous reports of increased GABA metabolism enzymes expression in hippocampus after ebselen treatment^70^. Given its synaptic location and dynamic regulation, *Gad2*, encoding the GAD65 isoform, is likely to play a critical role in linking metabolic with electrophysiological activity (Fig.5d-e). While it remains to be tested whether relative GAD conformation was altered and whether the rise in neurotransmitter cycling affected electrical activity in *Crtc1*^-/-^ mice, we speculate that the low Pi and PCr observed must create a shift from Apo- to Holo-GAD, which would drive a compensatory drop in mRNA level, as observed here, allowing this enzyme to maintain a stable rate of GABA synthesis (Fig.5g). This process would in turn favor the recycling of GABA for and from inhibitory neurotransmission rather than synthesis from glutamate, providing a mechanism to avoid excessive energy expenditure coming from extra metabolic steps, particularly when energy resources are low (Fig.6).

With the help of neuroimaging technologies such as MRS, MRI and PET we have identified potential clinically relevant biological markers with their associated environmental dependences, opening potential therapeutic strategies. Using high-field ^1^H-MRS we observed a drop in energy metabolites PCr and lactate in the hippocampus (Fig.1) that were associated with depressive-like behavior (Fig.3), suggesting their potential use as psychopathological ‘state’ markers. While both metabolites were found to be lower in *Crtc1*^-/-^ mice under basal conditions (i.e. 6 weeks of age; Fig1, Fig.3c and 4c) and associated with reduced glucose uptake measured with PET (Fig.2a-d), the addition of an external stressor (social isolation or OSFST) was able to modulate both the behavior and these *in vivo* markers (Fig.3 and 4). Stress, by challenging brain energetics, was shown to impact brain PCr content and behavior in chronic social defeat or chronic restraint protocols in mice^37,40^. Social isolation is known to affect the behavior in other rodents as well^71–73^ and induces several biological dysfunctions such as oxidative damage^74^, a loss of hippocampal parvalbumin neurons^75^ or drop in PCr content^76,77^. Accordingly, PCr and lactate levels appeared to relate tightly to the level of stress experienced and stimulating mitochondrial metabolism with ebselen was able to restore normal PCr levels together with normal behavior (Fig.4). Of note, *Crtc1*^-/-^ mice are very aggressive and show altered social behavior ^26^, thus rendering group housing more stressful for them than social isolation, which would explain the opposite behavioral and neurometabolic response observed compared to wild-type mice. Nevertheless, low lactate concentration, which can indicate both low glycolytic activity or high mitochondrial function, requires further considerations if it is to be indicative of a brain energetic status-based pathological state marker by itself. As such, hippocampal energy metabolite concentration showed a moderate ability to distinguish mice with ‘high’ and ‘low’ depressive-like behavior (Fig.S4f). Nevertheless, developing refined neuroimaging markers or functional paradigms to measure hippocampal neuroenergetics may allow significant clinical applications in the future. Furthermore, by combining MRI morphological analysis and ^1^H-MRS, we have identified putative inflammatory markers in the cingulate cortex (Fig.S5a-c, Fig.S2b and Fig.S3) that did not relate to the behavioral status, but reflected mouse genetic ‘susceptibility’. Specifically, we have been able to consistently observe an increase in prefrontal tCho, or specifically glycerophosphorylcholine (GPC) and phosphocholine (PCho), the degradation product and precursor of phosphatidylcholine (PtdCho) respectively, in *Crtc1*^-/-^ mice, together with PFC volume increase. Importantly, tCho concentration and tissue volume in PFC was able to differentiate *Crtc1*^-/-^ from wild-type mice (AUC: 82%), suggesting a potential use for clinical diagnosis or predicting treatment-compatibility. Interestingly, several MRS studies have reported elevated tCho levels in the anterior cingulate cortex of patients with bipolar disorders^78–81^ and these metabolites have been previously used as a MRS biomarker, such as for diagnostic of neoplastic tumor lesions in the brain^82–85^. Future research should address whether our observations in the DH and PFC could serve, respectively, as potential ‘diagnostic’ and ‘predictive’ clinical biomarkers for mood disorders. Finally, by identifying how *in vivo* brain markers associated with *Crtc1* respond to the environment, we provide a better characterization and understanding of the factors that influence the path from gene to depressive-like behavior, providing a hopeful step forward towards a precision medicine-based approach in the field of psychiatry.

## Methods

### Animals

*Crtc1* knock-out (*Crtc1*^-/-^) mice and wild-type (*Crtc1*^+/+^) littermates were bred and genotyped as previously described^26^. Mice were housed in standard Plexiglass filter-top cages in a normal 12h day-light cycle environment at a temperature of 23±1°C and humidity of 40%. Animals had *ad libitum* access to standard rodent chow diet and water. Weaning of newborn mice was done at 21 days and followed by group-housing until being isolated at ~6 weeks to prevent injuries of cage mates induced by the aggressive *Crtc1*^-/-^ male mice^26^. All experiments were carried out with the approval of the Cantonal Veterinary Authorities (Vaud, Switzerland) and conducted according to the Federal and Local ethical guidelines of Switzerland (Service de la consommation et des affaires vétérinaires, Epalinges, Switzerland) in compliance with the ARRIVE (Animal Research: Reporting *in vivo* Experiments) guidelines.

### Experimental design

Three sets of experimental designs of the present study in male *Crtc1*^-/-^ and wild-type (WT) mice were implemented. In the first experimental set, basal metabolic function was assessed and quantified in *Crtc1*^-/-^ and WT mice at the age of 6 weeks postnatal, and prior to the social separation from their littermates. The second experimental set was performed, longitudinally for 18 weeks, in mice aged from 6 to 24 weeks postnatal (Fig 3A). As before, the first measurement time point was at 6 weeks of age; thereafter, animals were socially isolated until the end of the study in an enriched environment that included a paper house and a wooden stick. The third set of experiments was conducted for 4 weeks, in mice aged from 6 to 10 weeks postnatal (Fig.4A). Animals were socially isolated again after the first imaging time point and were then subjected to a 4 weeks stress and treatment protocol.

### *In vivo* ^1^H-Magnetic Resonance Spectroscopy (^1^H-MRS)

Localized *in vivo* ^1^H-Magnetic Resonance Spectroscopy (^1^H-MRS) was performed in the dorsal hippocampus (DH) and cingulate prefrontal cortex (PFC) of *Crtc1*^-/-^ and wild-type mice. Animals were maintained under continuous isoflurane anesthesia (1.5% mixed with 1:1 air:oxygen mixture) and monitoring of physiology during the entire scan for physiological parameters. Breathing rate per minute was maintained between 70 – 100rpm using a small animal monitor (SA Instruments Inc., New York, USA) and rectal temperature was kept at 36.5±0.4°C with a circulating heating water bath and assessed using a temperature rectal probe. Animals were scanned in a horizontal 14.1T/26cm Varian magnet (Agilent Inc., USA) with a homemade transceiver ^1^H surface coil in quadrature. A set of T_2_-weighed images was acquired using a fast spin echo (FSE) sequence (15×0.4mm slices, TEeff/TR=50/2000ms, averages 2) to localize the volume of interest (VOI). The voxels were positioned to include either a single dorsal hippocampus (DH; 1×2×1 mm^3^) or the cingulate prefrontal cortex (PFC; 1.4×1.6×1.2 mm^3^). In each voxel, the field homogeneity was adjusted using FAST(EST)MAP^86^ to reach a typical water linewidth of 15±1Hz for DH and 14±1Hz for PFC. Proton spectra were acquired with a spin echo full intensity acquired localized (SPECIAL) sequence (TE/TR=2.8/4000ms)^87^ using VAPOR water suppression and outer volume suppression. Scans were acquired in blocks of 30 times 16 averages for DH and 8 averages for PFC. Post processing included frequency correction based on the creatine peak and summing of all the spectra before quantification with LCModel^88^. The water signal was used as internal reference and fitting quality was assessed using Cramer-Rao lower bounds errors (CRLB) for a typical rejection threshold of CRLB≥50%^89^. ^1^H-MRS acquisitions from DH and PFC of *Crtc1*^-/-^ mice and their wild-type littermates led to the reliable quantification of up to 20 individual metabolites, with a comparable spectra quality for both groups, i.e. with a signal-to-noise ratio (SNR) of 11.8±0.9 for wild-type vs. 13.4±1.3 for *Crtc1*^-/-^ in DH, and 15.0±0.7 for wild-type vs. 15.7±1.0 for *Crtc1*^-/-^ in PFC. MRI images acquired were used to quantify prefrontal cortex volume using a pattern-based morphometric approach. A surface in the shape of a kite was drawn on the coronal images with each corner situated between the major sulcus, the central lower part of the corpus callosum and the two cingulum bundles as reference points. The surface was quantified using ImageJ and averaged over the group for each brain section.

### High resolution NMR Spectroscopy

Mice were sacrificed using a microwave fixation apparatus (Gerling Applied Engineering Inc., Modesto, CA, USA) at 4kW for 0.6s after intraperitoneal injection of a lethal dose of sodium pentobarbital (~50μl to reach a dose of 150mg/kg). Brain was extracted, dorsal hippocampus was removed, frozen on liquid nitrogen and stored at -80°C. Samples were then ground on mortar using liquid nitrogen, weighed and followed by a CHCl_3_/MeOH Folch-Pi extraction^90,91^. Samples were stirred at 4°C in a 1:1:1 mixture of CHCl_3_:MeOH:H_2_O for 30min after what the aqueous phase was collected and lyophilized. The resulting extracted metabolites were resuspended in 600μl deuterium oxide containing 0.1μM 4,4-dimethyl-4-silapentane-1-sulfonic acid (DSS) as internal reference. High resolution NMR was performed using a DRX-600 spectrometer (Bruker BioSpin, Fällanden, Switzerland). Proton-NMR (^1^H-NMR) spectra were acquired with 400 scans using a pulse-acquired sequence (flip angle 30° and 5s pulse delay). Phosphorous-NMR (^31^P-NMR) spectra were acquired on the same sample with 10’000 scans using a proton-decoupled pulse-acquired sequence (flip angle 90° and 5s pulse delay). Spectra were analyzed and quantified using the MestReNova software (Mestrelab Research, Santiagio de Compostela, Spain). Spectra were phase and baseline corrected manually. Afterwards, peaks were integrated and referenced to the DSS resonance and normalized to NAA. NAA concentration in dorsal hippocampus was assumed to be 7mM as measured *in vivo*. The following resonance (δ, in ppm) were considered (number of protons, spectral pattern): AXP δ 6.13 (1H, d), creatine δ 3.026 (3H, s), phosphocreatine δ 3.028 (3H, s) and N-acetyl-aspartate δ 2.00 (1H, s). The following resonance were integrated in the ^31^P spectrum after setting the PCr resonance to 0 ppm: NAD^+^ δ -8.31 (2P, q), NADH δ -8.15 (2P, m), UDPGlc δ -9.83 (2P, m), Pi δ 3.8 (1P, m), GPC δ 3.07 (1P, s). The spectral pattern is described as follows: s, singlet; d, doublet; t, triplet; dd, doublet of doublet; m, multiplet. Due to overlap between resonances, the NADH/NAD^+^ ratio was calculated as follows: the left part of the NAD^+^ quadruplet (X=2.NADH+NAD^+^+UDPGlc) was integrated as well as the right part of the quadruplet (Y=NAD^+^) and the -9.83 ppm UDPGlc resonance (Z = UDPGlc). Then, NADH was obtained by subtracting Y and Z from X, followed by a division by 2. As we did not see changes in GPC in the hippocampus between groups, we used this signal as an internal reference for ^13^P-NMR spectra quantification.

### *In vivo* ^18^FDG positron emission tomography (^18^FDG-PET)

Dynamic non-invasive fluorodeoxyglucose positron emission tomography (^18^FDG-PET) was performed as described previously^59,92^. Briefly, mice under 1-2% (vol/vol) isoflurane anesthesia in O2 were positioned in the scanner after tail vein cannulation and remained monitored for temperature and breathing rate throughout the experiment. Imaging was performed after i.v. bolus injection of ^18^FDG (~50MBq) through the tail vein catheter within the first 20s of a 50 min duration PET scan. After histogramming and image reconstruction with the Labpet software (Gamma Medica, Sherbrook, Canada), PMOD 2.95 software (PMOD Technologies, Zurich) was used for the determination of the heat-maps of standardized uptake value (SUV), defined as (mean ROI activity [kBq/cm^3^])/(injected dose [kBq]/body weight [g]). Regions of interest, i.e. hippocampus (2×5.5 mm^2^), were manually drawn over one axial slice. Mathematical modeling of hippocampal glucose metabolism was performed as previously described^59,92^, using the radioactive decay-corrected activity density values in [kBq/cc]. Intergroup differences could not be attributed to differences in the amount of ^18^FDG entering the blood, body weight, nor to differences in the kinetics of the arterial input function.

### Gene expression analysis

Total RNA was extracted and purified from micropunches of dorsal hippocampus (DH) using a RNAeasy Plus Minikit (Qiagen, Venolo, Netherland) according to the manufacturer’s instructions. NanoDrop Lite (Thermo Scientific, Wilmington, DE, USA) was used for the UV spectrophotometric quantification of RNA concentrations and purity assessment. cDNAs were obtained by reverse transcription of the mRNA samples in 50μl reaction using Taqman Reagents and random hexamers (Applied Biosystems, Foster City, CA, USA). Real-time quantitative PCR was subsequently performed with cDNA concentrations of 0.16ng/μl on a 96-well plate with SYBR Green PCR Master Mix (Applied Biosystems). The reaction started with a 2min step at 50°C and 10min at 95°C, followed by 45 cycles of 15s at 95°C and 1min at 60°C. The relative gene expression was determined using the comparative ΔΔCt method and normalized to β-actine and β2 microglobulin (β-2m) as housekeeping genes. The primers were used at a concentration of 250nM and described in the Supplementary Table 1.

### Mitochondrial respirometry

Animals were sacrificed by rapid decapitation followed by dorsal hippocampus dissection. The tissue was weighed, placed in a petri dish on ice with 2ml of relaxing solution (2.8mM Ca2K_2_EGTA, 7.2mM K_2_EGTA, 5.8mM ATP, 6.6mM MgCl_2_, 20mM taurine, 15mM phosphocreatine, 20mM imidazole, 0.5mM dithiothreitol and 50mM MES, pH=7.1) until further preparation. Gentle homogenization was then performed in ice-cold respirometry medium (miR05: 0.5mM EGTA, 3mM MgCl_2_, 60mM potassium lactobionate, 20mM taurine, 10mM KH_2_PO_4_, 20mM HEPED, 110mM sucrose and 0.1% (w/v) BSA, pH=7.1) with an Eppendorf pestle. 2mg of tissue were then used for high resolution respirometry (Oroboros Oxygraph 2K, Oroboros Instruments, Innsbruck, Austria) to measure mitochondrial respiration rates at 37°C. The experimental protocol consists in several experimental steps, which test the capacity of the different mitochondrial electron transport chain components by measuring the O_2_ flux in the sample. 1) The activity of complex I (CI) is measured by adding ADP (5mM) to a mixture of malate (2mM), pyruvate (10mM) and glutamate (20mM). 2) Succinate (10mM) is subsequently added to the medium to stimulate complex II and measure the capacity of both complexes (CI+CII). 3) Protonophore FCCP (carbonyl cyanide 4-(trifluoromethoxy)phenylhydrazone) is then used (successive titrations of 0.2μM until reaching maximal respiration) to uncouple the respiration and provides information on the maximal capacity of the electron transfer system (ETS). 4) Rotenone (0.1μM) was then used to inhibit complex I and quantify the contribution of complex II in the uncoupled sate (ETS CII). 5) Antimycin (2μM) is added to inhibit complex III and block the ETS in order to assess the residual oxygen consumption (ROX) provided by oxidative reactions unrelated to mitochondrial respiration. Oxygen fluxes were normalized by the wet weight of tissue sample and corrected for ROX.

### Blood metabolite measurements

Blood sampling was performed after the last ^1^H-MRS scan of the longitudinal and treatment studies. Blood was collected from the trunk after head decapitation using collection tubes (Heparin/Li^+^ Microvette CB300 LH, Sarstedt). Samples were centrifugated at 1’000g for 10min at room temperature leading to ~100μL of plasma, which was then frozen in liquid nitrogen and stored at -80°C. Blood MeS markers were then quantified using an ELISA kit (insulin: EZRMI-13K, Millipore; glucose) and colorimetric assays (triglyceride: 10010303, Cayman;: 10009582, Cayman) according to the manufacturer’s instructions and with the following dilution factors: triglyceride: 1/2, insulin: 1/5, and glucose: 1/20.

### Open field test (OF)

The open-field test was used to assess mice locomotor activity^93^. Animals were placed in a white arena (50×50×40m^3^) illuminated with dimmed light (30lux). After 30min of habituation in the experiment room, mice were transferred to the center of the arena and were allowed to explore for 25min. Mice were tracked for 20min using a tracking software (Ethovision 11.0 XT, Noldus, Information Technology), after removing the habituation period of the 5 first minutes in each video. An analysis of these videos provided the mean distance travelled and mean velocity.

### Porsolt forced swim test (FST)

Animals were introduced into a 5L capacity cylinder of 15cm in diameter containing 23-25°C tap water in dimmed light (30lux) as described in Breuillaud et al.^26^. Water level in the cylinder was set to prevent the mouse from touching the bottom of the enclosure or to avoid any possible escape. The session was recorded with a camera positioned on top of the setup for 6min and videos were analyzed using a tracking software (Ethovision 11.0 XT, Noldus, Information Technology). Immobility time was measured after discarding the first minute of swimming in each video.

### Tail suspension test (TST)

Mice were suspended individually by the tail on a metal bar at a height of ~35cm. A stripe of adhesive tape was attached to the mouse tail at ~2cm from the extremity to perform the suspension to the bar. Animals were videotaped from the side of the setup and immobility time was recorded manually during 5min^26^.

### Composite behavior (averaged z-scores)

In the longitudinal study, a composite behavior was computed and considered both immobility times from the FST and TST reflecting animal’s behavioral despair. A z-score was calculated using MATLAB (Version 9.6, The MathsWorks Inc, Natick, MA) for each mouse and time-point using MATLAB function *normalize* with the option argument *zscore*. The z-score was computed using the overall average and standard deviation (including all mice and time-points). Finally, the behavioral composite z-score was calculated by averaging the two z-scores of TST and FST for each mouse time point.

### Repeated open-space forced swim test (OSFST)

The repeated OSFST protocol was used as described previously^26,27^. Animals were introduced into a cage (45×28×20cm) filled up to ~13cm with 34-35°C tap water colored with milk. Mice were subjected to 4 consecutive days of swimming (day -9 to -6) for 15min. Mice were then subjected to additional swim sessions for 3 weeks under treatment, according to the following interval: Days -1, 3, 7, 10, 13, 17, 20. Water was replaced regularly between tests to ensure constant water temperature. Animals were videotaped from above and immobility time was recorded manually.

### Ebselen treatment

Animals were treated with ebselen (Tokyo Chemical Industry, Tokyo, Japan) starting from day 0 until the end of the repeated OSFST protocol (Fig 4A). Mice received oral administration (gavage) of ebselen (10mg/kg) dissolved in 5% (w/v) carboxymethylcellulose (CMC; Sigma Aldrich) 2 times a day (mornings and evenings) for 21 consecutive days. The control group was administered a 5% CMC vehicle solution of the same volume. The dose was adjusted to any body weight gain.

### Neuroimaging marker assessment

Receiver operating characteristic (ROC) curves and the area under the ROC curve (AUC) were established for discriminating *Crtc1*^-/-^ from wild-type mice on the basis of their PFC status, which took into account the concentration of total choline (tCho) and tissue volume separately or as an average of individual z-scores. For this averaged score, the PFC individual z-scores were calculated using the whole sample average and standard deviation for both experiments combined (longitudinal and treatment). ROC curves and AUC were also established for the neuroenergetic profile (Lac and PCr) of DH for discriminating mice with ‘high’ and ‘low’ depressive-like behavior. In order to consider the different ways of assessing the behavior between the longitudinal (FST+TST) and treatment (OSFST) studies, a behavioral z-scores was calculated using the sample average and standard deviation for each behavioral test separately. Subsequently, mice were separated into ‘high’ or ‘low’ depressive-like behavior, whether their score was higher (+z) or lower (-z) than the average, respectively. The ability of the DH neuroimaging markers to distinguish these two populations was tested using ROC curves for either Lac or PCr concentrations separately or as an average of individual z-scores.

### *In vivo* indirect ^13^C Magnetic Resonance Spectroscopy (^1^H-[^13^C]-MRS)

Non-invasive indirect carbon-13 Magnetic Resonance Spectroscopy (^1^H-[^13^C]-MRS) was performed as previously described^59,94^. The experimental set-up was comparable to that of ^1^H-MRS, with two main differences: (1) animals underwent femoral vein cannulation for the infusion of uniformly labeled ^13^C-glucose ([U-^13^C_6_]Glc) for a scan of ~230min duration; and (2) the coil included a ^13^C channel. Breathing rate was maintained at ~80rpm and rectal body temperature was kept at 36.2±0.3°C for both groups throughout the scan. Blood glycemia was measured before (Glc_Blood_(WT)=7.7±3.5mM vs. Glc_Blood_(*Crtc1*^-/-^)=7.3±0.9mM, n.s.) and after the infusion/scan (Glc_Blood_(WT)=21±4mM vs. Glc_Blood_(*Crtc1*^-/-^)=28±13mM, n.s.) using a Breeze-2 meter (Bayer AG, Leverkusen, Germany). At the end of the experiment, blood lactate levels (Lacblood(WT)=7.7±1.0mM vs. Lacblood(*Crtc1*^-/-^)=7.9±0.9mM; n.s.) were measured using two nearby GM7 analyzers (Analox Instruments Ltd, Stourbridge, UK). The VOI included the bilateral dorsal hippocampus (2×5.5×1.5 mm^3^) and led to a typical water linewidth of 20±1Hz after field homogeneity adjustment. ^1^H-[^13^C]-MRS spectra were acquired using the full intensity SPECIAL-BISEP sequence (TE=2.8ms, TR=4000ms, averages=8) as previously described^59,95,96^. The non-edited (proton, ^1^H) and inverted spectra (editing OFF and ON) were obtained using an interleaved acquisition and were subtracted in the post processing steps to obtain the edited spectra (protons bound to carbon 13, ^1^H-[^13^C]). The non-edited spectra were quantified using a standard basis set for the neurochemical profile, while the edited spectra were fitted with a basis set that included simulated LacC3, LacC2, AlaC2+C3, GluC4, GluC3, GluC2, GlnC4, GlnC3, GlnC2, AspC3, AspC2, GABAC4, GABAC3, GABAC2 and acquired spectra of glucose. *In vivo* ^1^H-[^13^C]-MRS enables to follow the fate of brain glucose and its incorporation in several brain metabolites infusion of [U-^13^C_6_]Glc. Scanning the bilateral DH allowed us to quantify 12 metabolite resonances with a 10 min time resolution and a comparable SNR (as defined by the LCModel, i.e. the ratio of the maximum in the spectrum-minus-baseline to twice the rms residuals) between wild-type and *Crtc1*^-/-^ mice (SNR(^1^H):21.4±1.4 vs. 21.5±0.8; SNR(^1^H-[^13^C]):6.2±0.5 vs. 5.6±0.5, for wild-type and *Crtc1*^-/-^ respectively, mean±s.e.m). ^13^C concentration curves of each metabolite were determined by multiplying the fractional enrichment (FE) with the total molecular concentration measured in the non-edited spectra. Mathematical modeling was performed using either a “1-compartment” or a “pseudo 3-compartment” model of brain energy metabolism (see Cherix et al., 2020b for a complete description of the modeling). For both models, the cerebral metabolic rate of glucose (CMR_Glc_) was set to the value obtained in the same voxel from the ^18^FDG-PET experiments. Following fluxes were included in the 1-compartment model: tricarboxylic acid cycle (V_TCA_); a dilution flux from blood lactate (V_dil_^in^) and from blood acetate (V_dil_^g^); a transmitochondrial flux (V_x_); and finally, a neurotransmission flux (V_NT_). The estimated fluxes from the pseudo 3-compartment model (depicted on Fig.S5c) included: a dilution flux from blood lactate (V_dil_^in^) and from blood acetate (V_dil_^g^); the pyruvate dehydrogenase activity of excitatory (V_PDH_^e^) and inhibitory (V_PDH_^i^) neurons; a transmitochondrial flux for excitatory (V_x_^e^) and inhibitory (V_x_^i^) neurons; a neurotransmission flux for excitatory (V_NT_^e^) and inhibitory (V_NT_^i^) neurons; glutamate decarboxylase activity (VGAD); and two exchange fluxes between two Gln or two GABA pools (V_ex_^g^ and V_ex_^i^). Values of pyruvate carboxylase activity (V_PC_), glial tricarboxylic acid cycle (Vg) and glial transmitochondrial flux (V_x_^g^) were fixed to known values and glial Gln efflux (V_eff_) was set equal to V_PC_, as described in Cherix et al^59^. The other parameters were calculated from the estimated fluxes through mass-balance equations, assuming metabolic steady-state (i.e. no net change in metabolites concentration over the experiment duration): the GABA TCA shunt (V_shunt_^i^=VGAD-V_NT_^i^), glutamine synthetase activity (VGS=V_NT_^e^-V_NT_^i^+V_PC_), total GABA TCA (V_TCA_^i^ =V_PDH_^i^+V_shunt_^i^); total glial TCA (V_TCA_^g^ =Vg+V_PC_+V_NT_^i^), and the oxidative cerebral metabolic rate of glucose (CMR_Glc_(ox)=(V_TCA_^i^+V_TCA_^e^+V_TCA_^g^+V_PC_)/2). The brain-to-blood lactate efflux was calculated (V_dil_^out^= V_dil_^in^·Lacbrain/Lacblood) using the lactate concentration measured in the hippocampus (Lacbrain(WT)=2.5±1.1mM vs. Lacbrain(*Crtc1*^-/-^)=1.6±0.5mM; *P*<0.05, Student’s t-test), from the non-edited spectra quantification and the final blood lactate measurements (Lacblood). An allostatic load refers to an ‘excess’ in physiological/cellular dynamic adaption to match energetic needs in response to external stimuli^34^. To assess the level of mitochondrial allostatic pressure, the relative ‘oxidative allostatic loads’ for *Crtc1*^-/-^ mice were calculated for excitatory and inhibitory neurons separately, considering neurotransmission activity relative to mitochondrial ATP production, using following equation: Relative excitatory load = (V_NT_^e^/V_ATP(OX)_^e^)_Crtc1-/-_ / (V_NT_^e^/V_ATP(OX)_^e^)_WT_ and relative inhibitory load = ((V_NT_^i^+V_ex_^i^)/V_ATP(OX)_^i^)_Crtc1-/-_ / ((V_NT_^i^+V_ex_^i^)/V_ATP(OX)_^i^)_WT_, where V_NT_^e^ and (V_NT_^i^+V_ex_^i^) are the excitatory and inhibitory neurotransmission cycling activities respectively, and V_ATP(OX)_^e^ and V_ATP(OX)_^i^ are the excitatory and inhibitory ATP production rates from mitochondria (detailed in Fig.S5d).

### Statistics

Statistics were all performed with GraphPad Prism (GraphPad Software, San Diego, CA, USA). All values are given as mean±s.e.m. unless stated otherwise. P-values of *P*<0.05 were considered statistically significant. Metabolite data from high resolution ^1^H- and ^31^P-NMR were analyzed with a non-parametric Mann-Whitney test. Longitudinal measurements (behavior and metabolites) were analyzed using two-way analysis of variance (ANOVA) with genotype and time as both factors. Gene expression and metabolic comparisons with two factors (genotype and treatment) were analyzed with two-way ANOVA and a Bonferroni post-hoc test when appropriate. Data from the OSFST behavioral measurements were analyzed with a two-way ANOVA with repeated measures followed by a Fisher LSD post-hoc test^27^. Standard deviation of metabolic flux estimates was obtained from 300 Monte-Carlo simulations. Flux comparisons between *Crtc1*^-/-^ and wild-type mice were performed with a permutation analysis with 2000 random permutations, followed by individual two-tailed Student’s t-tests^97^. All the other comparisons between *Crtc1*^-/-^ and wild-type animals were performed with paired or unpaired Student t-test.

## Acknowledgements

This study was supported financially by the Center for Biomedical Imaging (CIBM) of the University of Lausanne (UNIL), University of Geneva (UNIGE), Geneva University Hospital (HUG), Lausanne University Hospital (CHUV), Swiss Federal Institute of Technology (EPFL) and the Leenaards and Louis-Jeantet Foundations and the Swiss National Science Foundation (Grants 31003A_149983 to RG and 31003A_170126 to JRC).

## Authors’ contributions

AC and JRC designed the study. AC, CPY, BL, OZ and JG acquired and analyzed the data. AC, CPY, CS, RG and JRC interpreted the data. AC drafted the manuscript. All the authors assisted in revising the manuscript and approved the final version.

## Ethics Declaration

The author(s) declare no potential conflicts of interest with respect to the research, authorship, and/or publication of this article.

## Supplementary Figures

**Figure S1:**
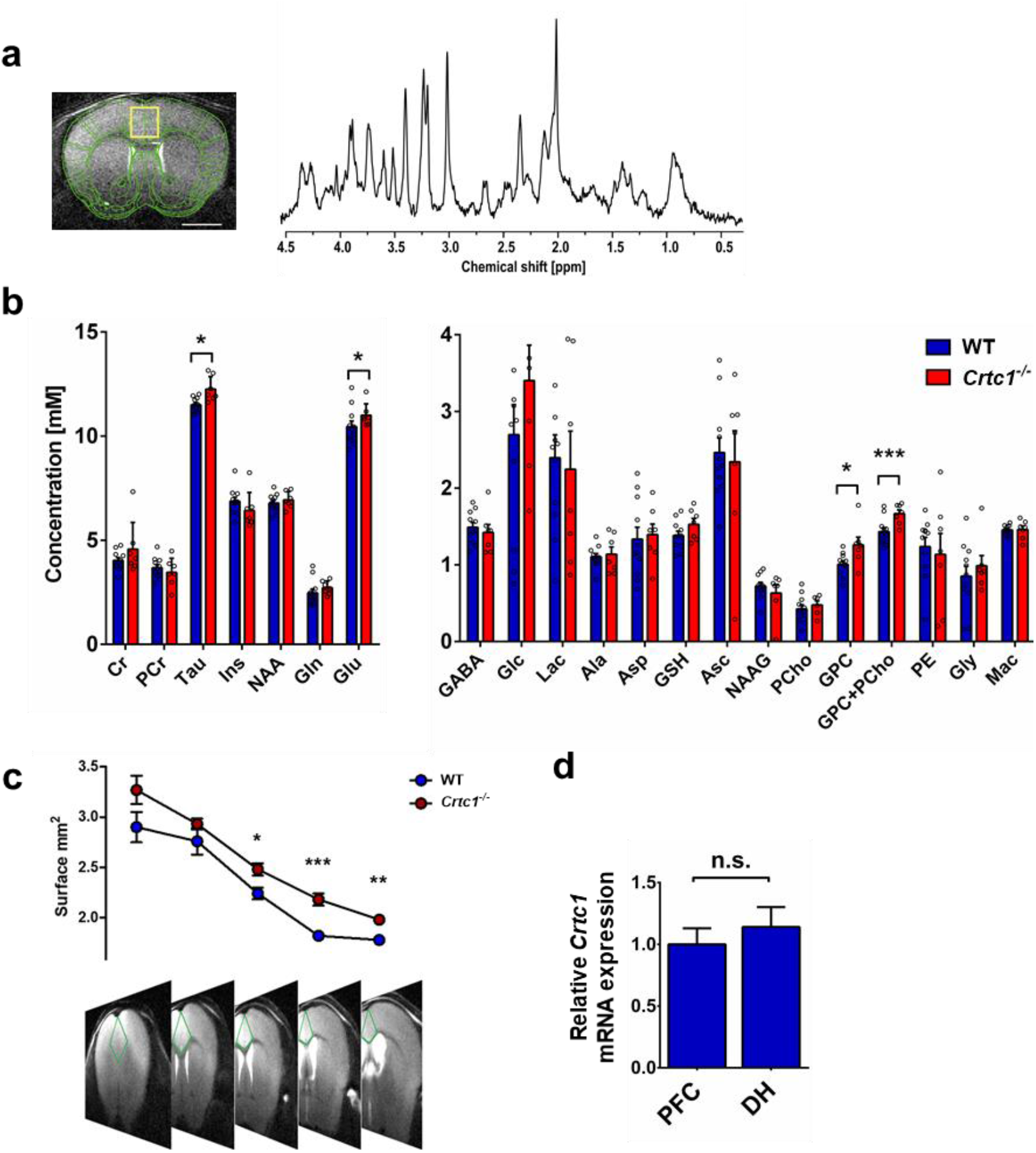
Deletion of *Crtc1* in mice is associated with increased inflammatory markers in PFC as revealed by MRS. **a**, T_2_-weighted image acquired for localized MRS (VOI including cingulate PFC: yellow rectangle) with a scale bar of 2mm (left) and typical ^1^H-MRS spectrum acquired in the VOI (right). **b**, Quantification of PFC neurochemical profile in 6 weeks old wild-type (WT; n=10) and *Crtc1*^-/-^ (n=6) mice, *** *P*<0.005, \**P*<0.05, unpaired Student’s t-test. **c**, Volumetric analysis of MRI images reveals a higher prefrontal volume in *Crtc1*^-/-^ as compared to wild-type mice (\*\*\**P*<0.005, ** *P*<0.01, * *P*<0.05, unpaired Student’s t-test). **d**, Relative *Crtc1* expression is not significantly different between PFC and dorsal hippocampus (DH) in wild-type mice (unpaired Student’s t-test, n.s., not significant). Data are shown as mean±s.e.m.

**Figure S2:**
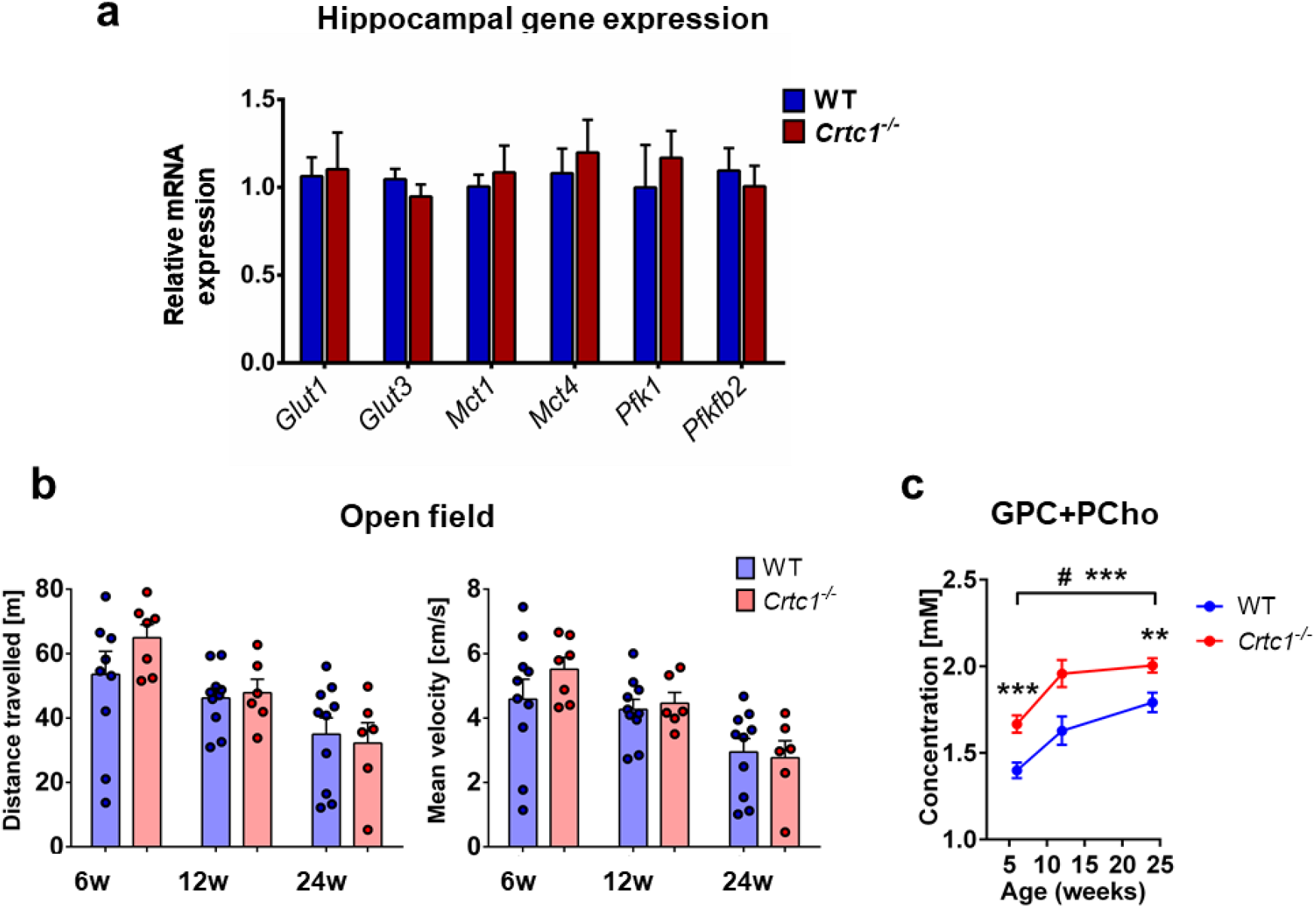
Supplementary data of the longitudinal experiment (6 to 24 weeks of age) **a**, Relative gene expression in hippocampus of wild-type (WT; n=10) and *Crtc1*^-/-^ (n=6) mice after the 18 weeks of social isolation. *Glut1-4*, Glucose transporter 1-4; *Mct1-4*, monocarboxylate transporter 1-4; *Pfk1*, phosphofructokinase 1; *Pfkfb2*, 6-phosphofructo-2-kinase/fructose-2,6-biphosphatase 2. **b**, Locomotor activity of wild-type (n=10) and *Crtc1*^-/-^ (n=6) mice during the longitudinal isolation experiment. **c**, Total choline (tCho; GPC+PCho) profile in PFC of wild-type (n=10) and *Crtc1*^-/-^ (n=6) mice during the isolation protocol. tCho remained increased in *Crtc1*^-/-^ independently of animal’s age (left panel; Genotype effect: F1,14=12.89, *P*<0.01; Time effect: F_2,28_=29.31, *P*<0.0001, two-way ANOVA, followed by Fisher LSD posthoc test; \*\*\**P*<0.005, \*\**P*<0.01, \**P*<0.05; # refers to wild-type only).

**Figure S3:**
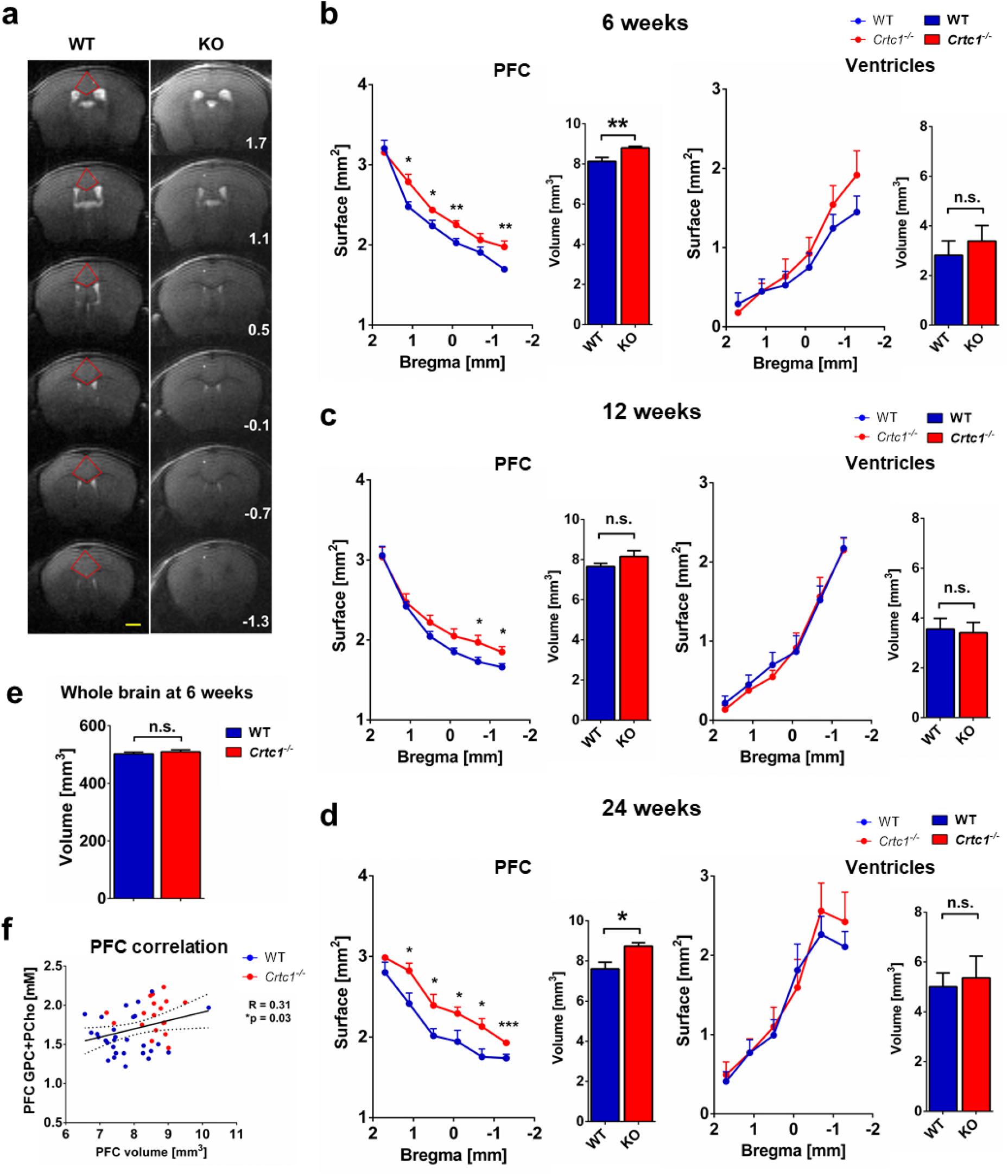
Volumetric analyses of prefrontal volume during the longitudinal experiment (6 to 24 weeks of age) **a**, Typical T_2_-weighted images analyzed with location of the ROI drawn on PFC (red kite); Yellow scale bar=2mm; white label: bregma coordinates (mm). **b-d**, Prefrontal (left panels) and ventricular (right panels) volumes measured from MRI images at the age of 6 weeks (b), 12 weeks (c) and 24 weeks (d). Unpaired Student’s t-test, \*\*\**P*<0.005, \*\**P*<0.01, \**P*<0.05, n.s., not significant. **e**, Whole brain volume measured from MRI images at the age of 6 weeks. **f**, Correlation between PFC total choline (tCho: GPC+PCho) and PFC volume (R=0.31, \**P*=0.03).

**Figure S4:**
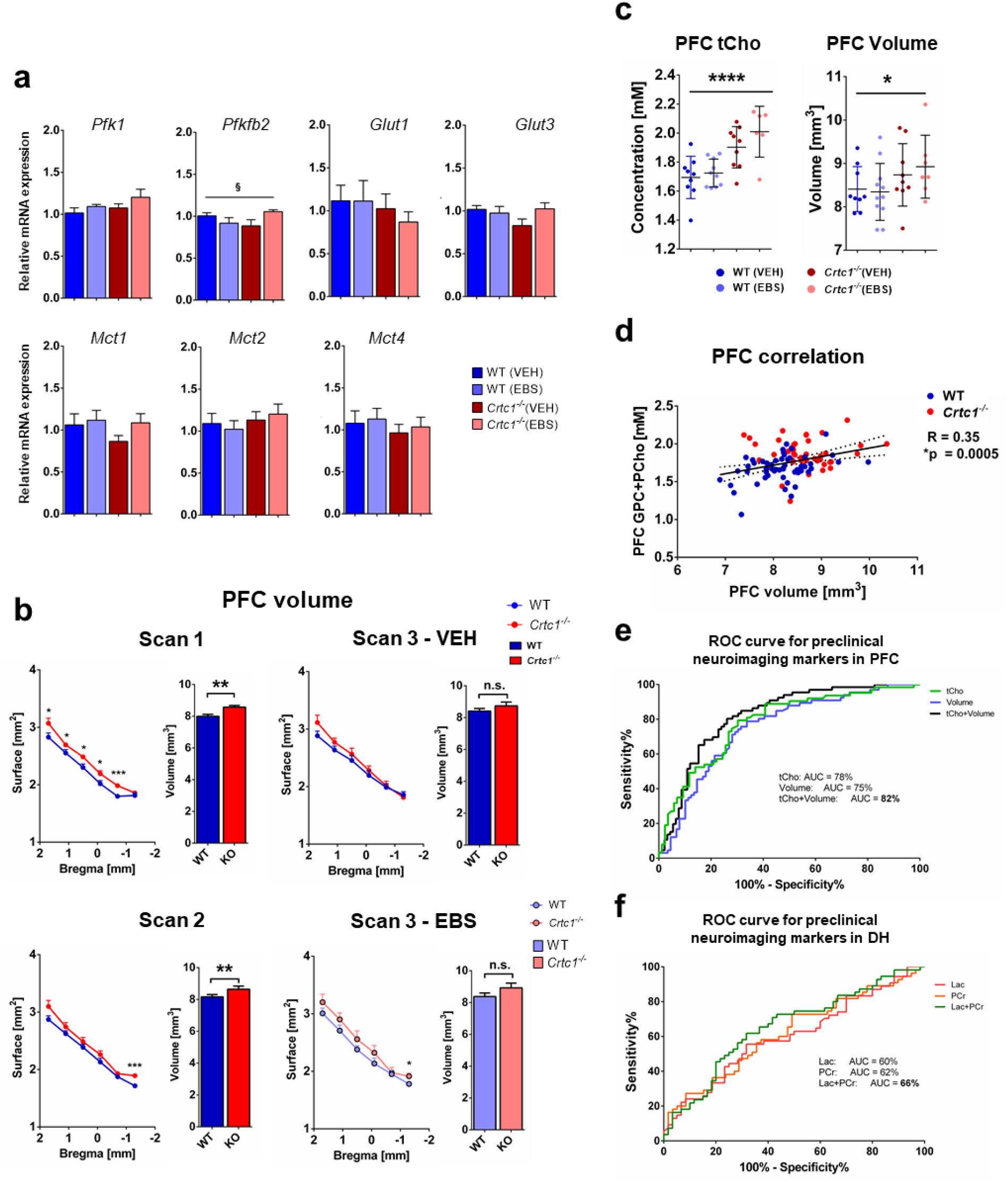
Supplementary hippocampal gene expression analyses and prefrontal volumetric analyses after ebselen treatment and OSFST. **a**, Hippocampal gene expression analysis after 21 days of ebselen treatment. (*Pfkfb2*: Interaction: F_1,28_=4.87, ^§^*P*=0.037, two-way ANOVA). *Pfk1*, phosphofructokinase 1; *Pfkfb2*, 6-phosphofructo-2-kinase/fructose-2,6-biphosphatase 2.; *Glut1-4*, Glucose transporter 1-4; *Mct1-4*, monocarboxylate transporter 1-4. **b,** Prefrontal volumetric analysis from neuroanatomical MRI images at baseline day - 10; Scan 1), before the start of the treatment (day -1; Scan 2) and at the end of the protocol (day 21) between the treated (EBS) and untreated (VEH) groups. Unpaired Student’s t-test, *** *P*<0.005, \*\**P*<0.01, * *P*<0.05, n.s., not significant. **c,** Prefrontal total choline concentration (tCho; GPC+PCho; left panel) at the end of the treatment (Genotype effect: F_1,28_=26.16, **** *P*<0.0001, two-way ANOVA). Prefrontal volume (right panel) at the end of the treatment (Genotype effect: F1,32=4.21, * *P*<0.05, two-way ANOVA). **d**, Correlation between PFC total choline (tCho: GPC+PCho) and PFC volume (R=0.35, \**P*=0.0005). **e**, Receiver operating characteristic (ROC) curves for prefrontal cortex (PFC) total choline (tCho) concentration (green), volume (blue) and the average of the z-scores of both measurements (tCho+Volume; black). Using a combination of both cholinergic and volumetric markers (averaged z-scores) provided an area under the curve (AUC) of 0.820 (95% CI 0.754-0.886), thus providing a good means of distinguishing *Crtc1*^-/-^ from wild-type mice. When considered separately, AUC for tCho was 0.783 (95% CI 0.709-0.858) and volume was 0.750 (95% CI 0.672-0.827). Analysis included samples from longitudinal (3 time points) and treatment (3 time points) studies for *Crtc1*^-/-^ (n=22) and wild-type (n=31) mice. **f**, ROC curves for energy metabolites lactate (Lac; red) and phosphocreatine (PCr, orange) and the average of the z-scores of both measurements (Lac+PCr; green) in dorsal hippocampus (DH) to distinguish mice with high versus low level of depressive-like behavior. Using a combination of both PCr and Lac provided an AUC of 0.656 (95% CI 0.555-0.756), while Lac had an AUC of 0.601 (95% CI 0.497-0.706) and PCr an AUC of 0.619 (95% CI 0.516-0.722), when considered separately. Analysis included samples from longitudinal (3 time points) and treatment (2 time points) studies for *Crtc1*^-/-^ (n=22) and wild-type (n=31) mice.

**Figure S5:**
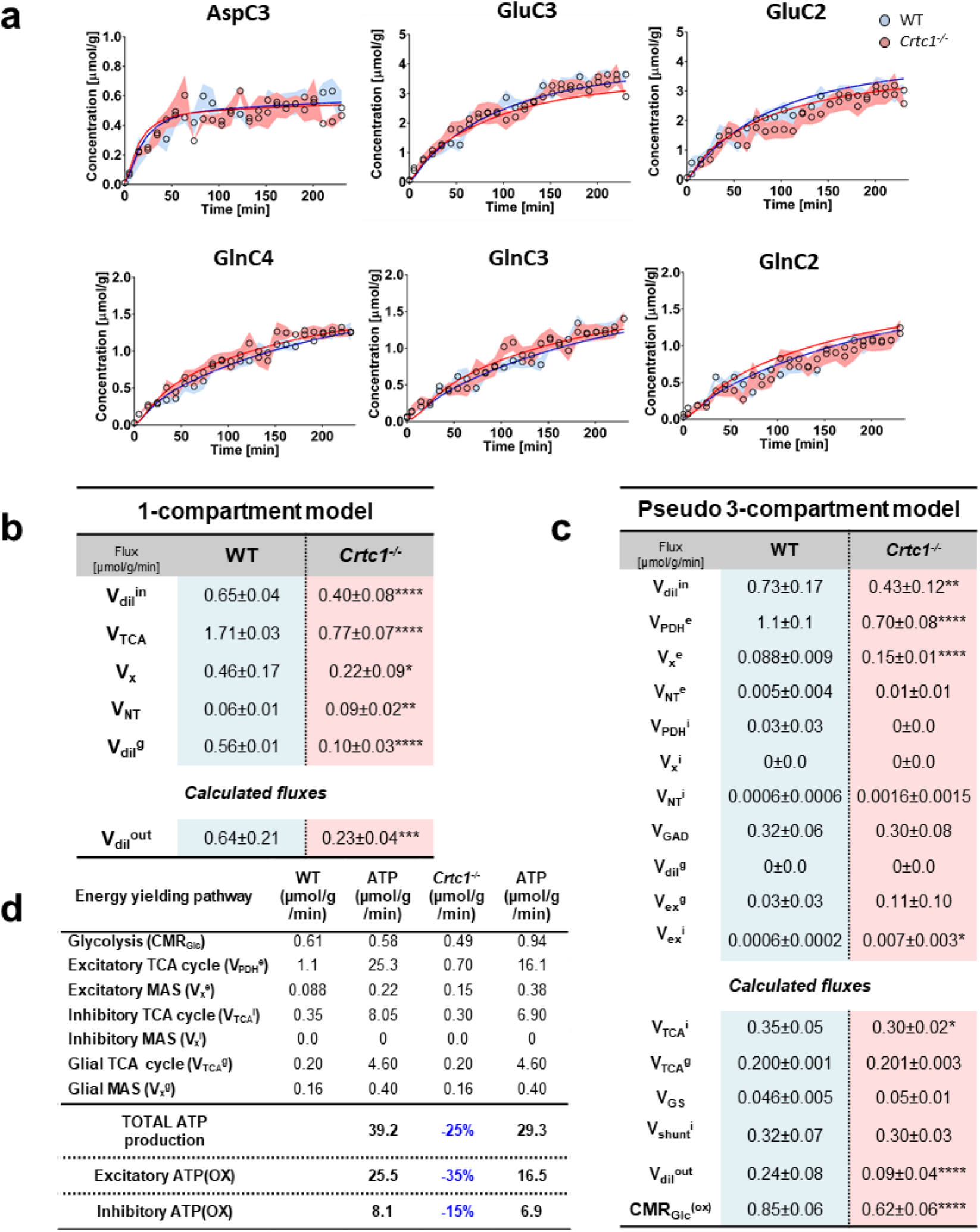
Supplementary ^13^C-labeling curves and metabolic flux values from mathematical modeling. **a**, Isotopic ^13^C-enrichment curves of remaining hippocampal metabolites included in the modeling (mean±s.d.) during ^1^H-[^13^C]-MRS experiment. Fitting of the data with a pseudo 3-compartment model of brain glucose metabolism is shown with a straight line for wild-type (WT; in blue) and *Crtc1*^-/-^ (in red) mice. **b-c**, Metabolic fluxes (mean±s.d.) determined using a pseudo 3-compartment model (b) or a 1-compartment model (c) of brain glucose metabolism. Estimated parameters from the 1-compartment model of brain energy metabolism: Blood lactate influx V_dil_^in^; TCA cycle V_TCA_; transmitochondrial flux V_x_; neurotransmission flux V_NT_ and glial dilution factor V_dil_^g^. Estimated parameters from the pseudo 3-compartment model of brain energy metabolism (fluxes are separated into glutamatergic (^e^), GABAergic (^i^) and glial (^g^) compartments): The pyruvate dehydrogenase activity (V_PDH_), glial tricarboxylic acid cycle (Vg), a dilution flux from blood lactate (V_dil_^in^) and from blood acetate (V_dil_^g^), a transmitochondrial flux (V_x_), a neurotransmission flux (V_NT_), pyruvate carboxylase flux (V_PC_), a Gln efflux (V_eff_), glutamine synthetase activity (V_GS_), glutamate decarboxylase activity (VGAD), GABA TCA shunt (V_shunt_) and two exchange fluxes between two Gln or two GABA pools (V_ex_^g^ and V_ex_^i^). Parameters calculated from these metabolic fluxes: inhibitory TCA cycle V_TCA_^i^; glial TCA cycle rate V_TCA_^g^; glutamine synthetase activity VGS; GABA shunt rate V_shunt_^i^(=V_shunt_^g^); lactate blood efflux V_dil_^out^; total TCA cycle or the oxidative cerebral metabolic rate of glucose CMR_Glc_(ox). Cerebral metabolic rate of glucose (CMR_Glc_) was used from the ^18^FDG-PET experiment. All the *P* values are from unpaired Student t-test, \**P*<0.05, \*\**P*<0.005, \*\*\**P*<0.0005, \*\*\*\**P*<0.0001. Flux estimates are reported with the standard deviation generated by the MC simulation. All fluxes are given in μmol/g/min. **d,** The ATP production (in μmol/g/min) was calculated with known yields (from Hertz et al^98^.) for each energy pathway and compared between excitatory and inhibitory contributions. Consumption of one molecule of glucose produces 2 ATP and 2 NADH. Mitochondrial function produces ~23 ATP from the action of pyruvate dehydrogenase (PDH) and tricarboxylic acid (TCA) per pyruvate molecule. The malate-aspartate shuttle (MAS) fuels mitochondrial electron transport system (ETS) with cytoplasmic reducing equivalents, yielding ~2.5 ATP per molecule of NADH.

**Table S1:**
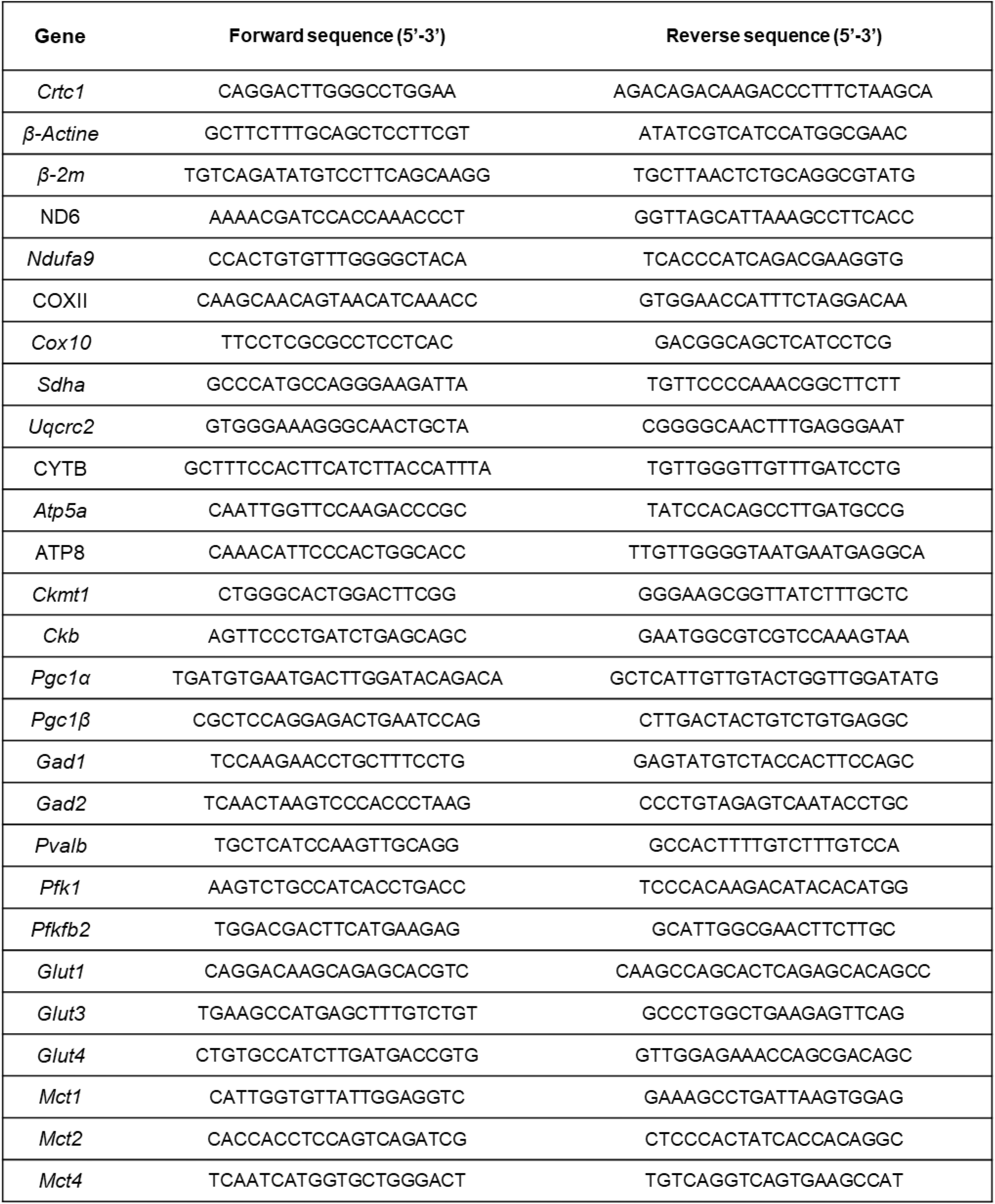
List of primers. *Crtc1*: CREB Regulated Transcription Coactivator 1; β-2m: β2 microglobulin; ND6: Mitochondrially Encoded NADH:Ubiquinone Oxidoreductase Core Subunit 6; *Ndufa9:* NADH:ubiquinone oxidoreductase subunit A9; COXII: Cytochrome c oxidase subunit 2; *Cox10*: Cytochrome C Oxidase Assembly Factor Heme A:Farnesyltransferase COX10; *Sdha*: Succinate Dehydrogenase Complex Flavoprotein Subunit A; *Uqcrc2:* Ubiquinol-Cytochrome C Reductase Core Protein 2; CYTB: mitochondrially encoded cytochrome b; *Atp5a:* ATP synthase lipid-binding protein; ATP8: Mitochondrially Encoded ATP Synthase Membrane Subunit 8; *Ckmt1:* Creatine kinase U-type, mitochondrial; *Ckb:* Creatine kinase B-type; *Pgc1α:* Peroxisome Proliferator-Activated Receptor Gamma Coactivator 1-Alpha; *Pgc1* β: Peroxisome Proliferator-Activated Receptor Gamma Coactivator 1-Beta; *Gad1:* glutamate decarboxylase 1; *Gad2:* glutamate decarboxylase 2; *Pvalb:* Parvalbumin; *Pfk1:* ATP-dependent 6-phosphofructokinase subunit alpha; *Pfkfb2:* 6-phosphofructo-2-kinase/fructose-2,6-bisphosphatase 2; *Glut1-4:* facilitated glucose transporter, member 1-4; *Mct1-4:* monocarboxylic acid transporters, member 1-4.

